# Chemogenetic activation of hippocampal area CA2 promotes acute and chronic seizures in a mouse model of epilepsy

**DOI:** 10.64898/2025.12.29.696896

**Authors:** John J. LaFrancois, Meghan Kennedy, Monarchsinh Rathod, Bina Santoro, Christos Panagiotis Lisgaras, Steven A. Siegelbaum, Helen E. Scharfman

## Abstract

Pyramidal cells (PCs) of hippocampal area CA2 exhibit increased excitability in temporal lobe epilepsy (TLE) and in mouse models of TLE. In epileptic mice, selective inhibition of CA2 PCs reduces chronic seizures. Here we asked if activating CA2 PCs increases seizures. Mice expressing Cre recombinase in CA2 PCs (*Amigo2*-Cre mice) were injected with the convulsant pilocarpine to induce a period of severe seizures (*status epilepticus*, SE), which leads to chronic seizures after 3-4 weeks (epilepsy). Epileptic mice were injected with a Cre-dependent adeno-associated virus (AAV) to express an excitatory designer receptor exclusively activated by designer drug (eDREADD; hM3Dq) in dorsal CA2 bilaterally and implanted with subdural EEG electrodes. After recovery, mice were recorded continuously using video and EEG for 6 weeks, 3 weeks with drinking water containing the eDREADD activator clozapine-N-oxide (CNO) and 3 weeks without CNO. CA2 activation with CNO caused a significant increase in seizure frequency and duration. Seizures occurred in clusters (many seizures per day over several consecutive days) and mice given water with CNO had a greater maximum number of seizures per day during a cluster compared to water without CNO. CNO had no significant effect in control mice. In naïve *Amigo2-*Cre mice expressing hM3Dq, pre-treatment with CNO before pilocarpine administration shortened the latency to SE and increased EEG power at the start of SE. Taken together with prior findings, the results suggest that CA2 is a control point for regulating seizures in the pilocarpine mouse model of TLE.

**HIGHLIGHTS:**

- Chemogenetic excitation of CA2 increased chronic seizure frequency and duration in epileptic mice.
- When seizures occurred in clusters, CA2 excitation increased the peak of the cluster.
- When sexes were separated, males showed effects on seizure frequency but both sexes showed effects on seizure duration and clusters.
- Chemogenetic excitation of CA2 promoted pilocarpine-induced *status epilepticus* in normal mice.

**GRAPHICAL ABSTRACT:** Legend: In adult mice, injecting the convulsant pilocarpine induces severe seizures (*status epilepticus*, SE) and ultimately leads to chronic intermittent seizures (epilepsy). Selective activation of hippocampal area CA2 during the period of chronic seizures led to greater seizure frequency and duration, as well as a greater number of seizures/day at the peak of a seizure cluster. Effects were greatest in males. In naive mice, activation of CA2 prior to pilocarpine administration facilitated SE. Therefore, CA2 influences acute and chronic seizures in this mouse model.

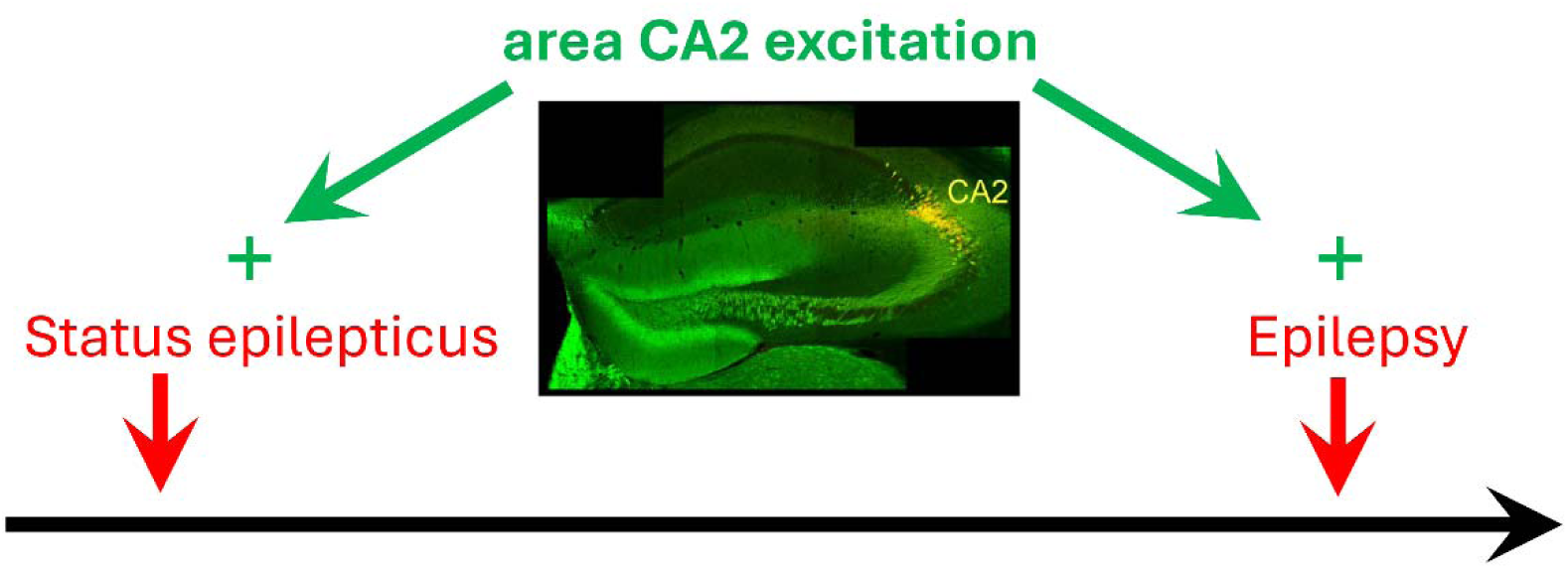

## INTRODUCTION

Several studies have implicated area CA2 of hippocampus in epilepsy. These investigations were initially motivated by observations that CA2 was relatively resistant to the pathology found in temporal lobe epilepsy (TLE) in humans, originally called hippocampal sclerosis (Margerison and Corsellis, 1966; Falconer, 1974; Bruton, 1988; Sloviter, 1989; Leranth and Ribak, 1991; Blümcke et al., 2012). Once rat models of TLE became available, resistance of area CA2 to neuronal loss was found in the rodents also (Meldrum, 1986; Sloviter, 1989; Sperk, 1994). Subsequent studies in mice found similar resistance of CA2 (Winawer et al., 2007; Whitebirch et al., 2022; Kilias et al., 2023). Importantly, when area CA2 was examined electrophysiologically in hippocampal slices of patients with medically refractory TLE, in which the hippocampus was removed to reduce seizures, CA2 neurons exhibited spontaneous bursts of action potentials, suggesting they were hyperexcitable in TLE compared to normal conditions (Wittner et al., 2009; Reyes-Garcia et al., 2018). An area corresponding to CA2 and CA3a also has spontaneous bursts of neuronal activity in slices from normal rodents treated with convulsants (Miles et al., 1984; Voskuyl and Albus, 1985; Tancredi and Avoli, 1987).

After transgenic mice with Cre recombinase expression in CA2 pyramidal cells (PCs) were generated (Hitti and Siegelbaum, 2014), and designer receptors exclusively activated by designer drugs (DREADDs) were developed (Urban and Roth, 2015), it became possible to inhibit CA2 selectively using an agonist to activate the inhibitory DREADD hM4Di in a common mouse model of TLE (Whitebirch et al., 2022). This model uses the cholinergic muscarinic agonist pilocarpine to induce acute severe seizures, known as *status epilepticus* (SE), which leads to lifelong chronic seizures (epilepsy) after several weeks (Turski et al., 1984). Using *Amigo2*-Cre epileptic mice, it was shown that DREADD inhibition of CA2 PCs reduced chronic seizures (Whitebirch et al., 2022). However, the converse was never tested, i.e., whether excitation of CA2 PCs would increase chronic seizures. This question is important because it is possible excitation would not increase seizures since CA2 PCs already are hyperexcitable in the epileptic brain (Wittner et al., 2009; Whitebirch et al., 2022), so that any additional excitation might have little effect. In addition, previous studies have highlighted a complexity in the ways CA2 influences adjacent hippocampal regions. For example, data from chronic silencing of CA2 using tetanus toxin light chain revealed an inhibitory effect of CA2 on CA3 recurrent excitatory collaterals (Boehringer et al., 2017). Therefore, if one were to use chemogenetics to activate CA2 it might depress the CA3 recurrent collateral pathway further. Since the CA3 recurrent collaterals are thought to be critical to the generation of synchronized epileptiform discharges and seizures (Alexander et al., 2018; Ren et al., 2021), the net effect might be to reduce seizures instead of promoting them. These complexities further supported our study of CA2 in epileptic mice using chemogenetic excitation.

To test the effects of CA2 excitation, *Amigo2*-Cre mice were injected with pilocarpine to induce SE. After epilepsy was established, i.e., after spontaneous intermittent seizures developed, mice were injected bilaterally in dorsal hippocampus with a Cre-dependent virus to express the eDREADD hM3Dq tagged with the fluorescent marker mCherry in CA2. After recovery and viral expression, the mice were recorded by video and EEG continuously and given water containing the eDREADD activator clozapine-N-oxide (CNO) for 3 weeks and water without CNO for 3 weeks. In addition, mice were also examined to determine if chemogenetic excitation of CA2 prior to SE would modify SE.

The results show that activating area CA2 during chronic epilepsy increases seizure frequency and seizure duration. Clusters of seizures were more severe. The effects on seizure frequency were detected only in males but both sexes showed effects on seizure duration and seizure clusters. In studies of SE, activation of area CA2 shortened the latency to SE and increased power during the initial phase of SE. Together with prior findings (Whitebirch et al., 2022), the results support a bidirectional regulation of CA2 in the pilocarpine model of epilepsy.

## METHODS

### Animals and animal care

All procedures were reviewed and approved by the Institutional Animal Care and Use Committee of The Nathan Kline Institute. Experiments were performed in accordance with guidelines of the Nathan Kline Institute and national guidelines for the care and use of laboratory animals.

We used the *Amigo2*-Cre mouse line that enables Cre-dependent AAV expression localized to CA2 PCs in hippocampus based on co-expression with CA2 markers (Purkinje cell protein 4; PCP4, Regulator of G-protein signaling 14; RGS14; (Laeremans et al., 2013; Evans et al., 2014; Hitti and Siegelbaum, 2014; San Antonio et al., 2014; Radzicki et al., 2023). *Amigo2*-Cre^+/-^ males were backcrossed to C57BL/6NCrl females for >10 generations (Cat#027, Charles River Laboratories). Animals used for experiments were produced by breeding *Amigo2*-Cre^+/-^ males to C57BL/6NCrl females. For breeding, mothers were provided a nestlet (2”x2”; W.F. Fisher) and fed Purina 5008 chow (W.F. Fisher). After weaning at 25-30 days of age, mice were housed with others of the same sex (2-4/cage) and fed Purina 5001 chow (W.F. Fisher) *ad libitum*. They were housed in standard mouse cages with a 12 hour light:dark cycle (lights on: 7:00 a.m.-7:00 p.m.) and relative humidity between 64-78%. Animals underwent procedures as soon as they reached the appropriate age (i.e., 8 weeks-old), independent of genotype. They were assigned to experimental or control groups randomly. Data were analyzed as soon as they became available, independent of the group. This approach was used to randomize data acquisition and analysis. After surgery to implant EEG electrodes, mice were housed alone because cage mates often scratched a wound area. Tail samples were used for genotyping (Mouse Genotyping Core, New York University Medical Center).

### Stereotaxic injection of viruses and EEG electrodes

Mice were approximately 8 weeks-old at the time of surgery. Prior to surgery, the analgesic Buprenex (Buprenorphine hydrochloride; NDC# 1296-0757-5; Reckitt Benckheiser) was diluted in sterile saline to yield a 0.03 mg/ml stock solution and 0.2 mg/kg was injected s.c. Mice were anesthetized by 3% isoflurane Aerrane, Patterson Veterinary) by inhalation in an induction chamber (#050XS; Kent Scientific), then transferred to a rodent stereotaxic apparatus (#902, David Kopf). Body temperature was maintained at 37°C with a homeothermic heating pad (Somnosuite, Kent Scientific). Isoflurane (1-2%, mixed with oxygen) was delivered through a nose cone (World Precision Instruments, WPI) to maintain anesthesia during surgery. The skin of the head was shaved, sterilized with Betadine (Purdue Products) and 70% ethanol (Pharmaco-Aaper), and then a midline incision was made in the skin over the skull. Fascia was cleared and a hand drill (#177001, Dremel Instruments) was used to create a craniotomy above the injection sites.

Using an automated syringe pump (#UMP3, WPI) with a 500 nl syringe (#7000.5, Hamilton) and a 25-gauge needle, 200-230 nl of AAV-hSyn-DIO-hM3Dq-mCherry (Addgene 44361-AAV2) or AAV-hSyn-DIO-mCherry (Addgene 50459-AAV2) was injected into hippocampus at 40 nl/minute. Titers for all injected viruses were ∼10^12^ vector genomes (vg) per ml. Aliquots of 5 µl of each virus were stored at −80°C until the day of use. The coordinates (in mm from Bregma) for injections were: Anterior-Posterior (A-P) 1.7, Medio-Lateral (M-L) 1.7, Dorso-Ventral (D-V) 1.7 relative to the skull surface. The needle was left at the injection depth for 5 minutes and then raised to half the depth for another 5 minutes to allow diffusion of the virus before being slowly retracted. Next, 0.10 inch-long subdural screw electrodes (stainless steel jeweler’s screws, #P000120CE094, J.I. Morris) were secured in the 2 craniotomies. Craniotomies were also made for additional subdural screw electrodes over the right occipital cortex (A-P - 3.5, M-L, +2.0), left frontal cortex (A-P −0.4, M-L −1.5), right olfactory bulb as ground (A-P +2.3, M-L +1.8), and the cerebellum as reference (A-P −5.7, M-L −0.5). For all electrode locations, the coordinates were adjusted proportionally if the distance between Bregma and Lambda was less than the standard for adult mice, 4.2 mm (Paxinos et al., 2001). Electrodes were attached to an 8-pin connector that was centered over the skull and secured with dental cement. Animals were transferred to a clean cage on a heating pad to keep the cage temperature above 30°C until they recovered from anesthesia. They were housed in the room where recordings would be made so they could acclimate to the environment.

### Induction of SE

AAV was injected and electrodes were implanted at approximately 8 weeks of age as described above. Three weeks later, pilocarpine was administered. Baseline recordings before drug injection were acquired for at least 1 hour. Following the baseline period, mice were injected subcutaneously (s.c.) with the peripheral muscarinic antagonist scopolamine methyl nitrate (1 mg/kg of 0.2 mg/ml in sterile 0.9% sodium chloride solution; Cat# S2250, Millipore Sigma) to reduce the peripheral effects of pilocarpine. The β2-adrenergic agonist terbutaline hemisulfate (1 mg/kg of 0.2 mg/ml in sterile 0.9% sodium chloride solution; Cat# T2528, Millipore Sigma) was administered s.c. with scopolamine to support respiration (Cho et al., 2015). The antiseizure drug ethosuximide (150 mg/kg of 84 mg/ml in phosphate buffered saline; Cat# E;7138, Millipore Sigma) was administered s.c. as a separate injection because it precipitated with the other solution when combined. Ethosuximide was used because it reduces the severe seizures during SE that cause death (Iyengar et al., 2015). Thirty minutes later, mice were injected s.c. with pilocarpine hydrochloride (210-230 mg/kg of 50 mg/ml in sterile 0.9% sodium chloride solution s.c. (#P6503, Millipore Sigma). Different doses were used because different batches of pilocarpine had different potency. All mice were injected s.c. with diazepam (10 mg/kg of 5 mg/ml stock solution; NDC# 0409-3213-12, Hospira) 2 hours after the pilocarpine injection to reduce the severity of SE, which improves morbidity and mortality after SE (Goodkin and Kapur, 2009; Iyengar et al., 2015). While sedated with diazepam, animals were injected with 2.5 ml warm (31°C) lactated Ringer’s solution s.c. (NDC# 07-893-1389, Aspen Veterinary Resources). Another injection was given approximately 4 hours later, and again 1-2x the next day. For the next 3 days, food was placed at the base of the cage and supplemented with (DietGel76A; ClearH_2_O). The cage was placed on a heating blanket to maintain cage temperature above 30°C.

### Video-EEG (vEEG)

#### Chronic epilepsy

Mice were allowed 4 weeks to recover from AAV injection and EEG electrode implantation. During this time, mice were housed in the room where vEEG equipment is located so that mice could acclimate to the recording environment. To record vEEG, the pin connector on the head of the mouse was attached to a preamplifier (Cat# 8406, Pinnacle Technology) which was attached to a mouse swivel/commutator (Cat# 8408, Pinnacle Technology) to allow freedom of movement. Signals were acquired at a 500 Hz sampling rate, and band-pass filtered at 1-100 Hz using Sirenia Acquisition software (https://www.pinnaclet.com, RRID:SCR_016183). Video was captured with a high-intensity infrared light-emitting diode (LED) camera (#1AC-2612C-BS, Zosi) and was synchronized to the EEG record. Mice were recorded 24 hours/day for 6 weeks or 9 weeks (Figure 1A).

**Figure 1.**
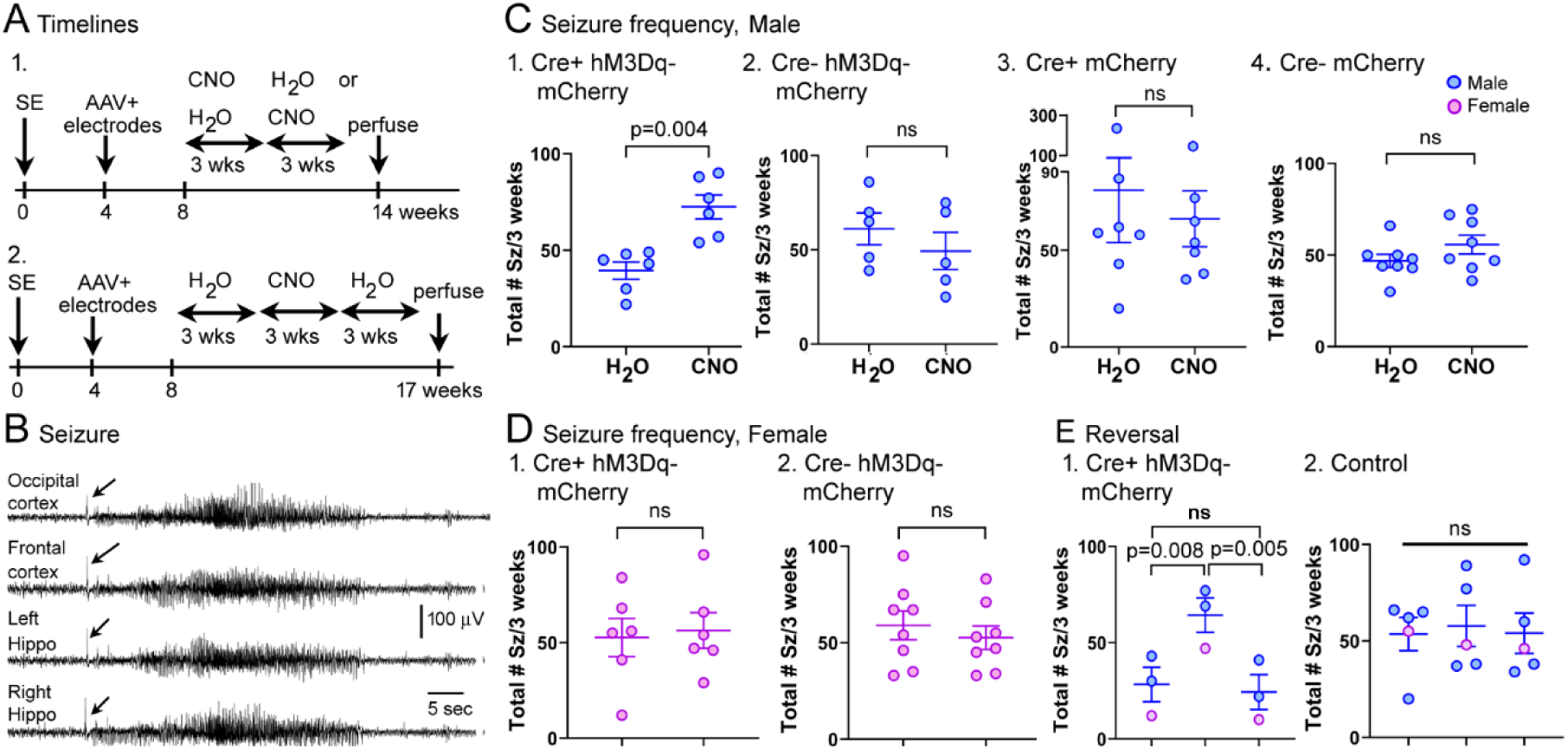
Chemogenetic excitation of CA2 in *Amigo2*-Cre+ mice increases seizure frequency. **A.** Experimental design.

1. SE was induced at approximately 8 weeks of age (time 0). AAV was injected and electrodes were implanted 4 weeks later. Four weeks after surgery mice were recorded continuously with video and EEG (vEEG) and provided water with CNO or water without CNO for 3 weeks. Then mice were provided the opposite treatment for another 3 weeks. Subsequently mice were perfused.
2. In a subset of mice there was an additional 3 weeks of continuous vEEG to determine if there was reversal of the effects of CNO. These mice received water without CNO, then water with CNO, and then an additional 3 weeks of water without CNO.
**B.** Example of a typical stage 5 convulsive seizure. The recordings from each electrode are shown. The arrows point to sentinel spikes that mark the onset of the seizure.
**C.** The total number of seizures for 3 weeks is shown for 4 groups of males. CNO increased the number of seizures in male Cre+ mice injected with AAV-hM3Dq-mCherry (C1, n=6 mice) but had no significant effect in the 3 control groups (C2, n=5; C3, n=7; C4, n=8). ns=not significant. Statistical comparisons were paired t-tests for C-D and are shown in Supplemental Table 1.
**D.** Seizure frequency in females showed no significant effects of CNO (D1, n=6; D2, n=8).

1. Mice were treated with water that had no CNO for 3 weeks followed by water with CNO for 3 weeks and finally an additional 3 weeks of water without CNO. There was a difference in the treatments in Cre+ mice (RM one-way ANOVA, F(2,4)=28.00, p=0.004; n=2 males, 1 female) with a significant increase in seizures from the first 3 weeks with water that had no CNO compared to the 3 weeks with water containing CNO (Tukey *post-hoc* test, p=0.008). There was a significant decline between the 3 week treatment with CNO and return to water without CNO (p=0.005), but no difference between the initial and last water treatments (p=0.787).
2. In controls expressing mCherry, seizures were not significantly different between treatments (E2, RM one-way ANOVA, F(2,8)=0.23, p=0.799, n=3 Cre+ males injected with AAV-mCherry and 2 Cre-mice injected with mCherry, 1 male and 1 female).

#### Acute seizures

AAV was injected and electrodes were implanted at approximately 8 weeks of age as described above. Three weeks later, pilocarpine was administered. Video-EEG was recorded for an hour before and 72 hours after pilocarpine injection.

### Chemogenetics

To address potential effects of hM3Dq on chronic seizures, mice were recorded continuously for 3 weeks while drinking water with CNO (10 mg/kg/day) and another 3 weeks with water that had no CNO. Some animals received water without CNO first and others received water with CNO first. The order was randomized. In a separate cohort, mice received water without CNO, then water with CNO, and then water without CNO (3 weeks each). CNO was prepared as a stock solution (1 mg/ml dissolved in sterile saline) stored at 4°C, and a fresh stock solution was made every 7 days. Stock solution was diluted in ddH_2_O to provide a dose of 10mg/kg/day based on a water consumption of 6 ml per day (described below). Water was changed every Tuesday and Friday.

To determine if mice drank a different amount of water when CNO was included, comparisons were made of water consumption. To determine the volume of water mice consumed per day, 5 mice were studied (2 male Cre+, 1 male Cre-, 1 female Cre+, 1 female Cre). These mice had SE, were implanted with electrodes, and had viral injections. There was only one mouse per cage. Volumes of water were measured in a graduated cylinder. Measurements were replicated each week for the 6 weeks of treatment (3 weeks with CNO in the water, 3 weeks without CNO in the water, order randomized). Supplemental Figure 1 shows that the mean volume consumed did not differ significantly between water with and without CNO (paired t-test, t=0.40, df=4, p=0.712). The mean volume per day was approximately 6 ml, which was also calculated in a previous study (Whitebirch et al.., 2022).

For analysis of the effects of hM3Dq on acute seizures during SE, 10 mg/kg CNO (dissolved in sterile saline, s.c.) was injected 30 minutes before pilocarpine in Cre+ and Cre-mice.

### Seizure analysis

#### Chronic epilepsy

EEG was analyzed offline with Sirenia Seizure Pro (V2.0.7; Pinnacle Technology, RRID:SCR_016184). A seizure was defined as a period of rhythmic (>3 Hz) deflections that were >2x the standard deviation of mean peak-to-peak amplitude of a 10 second period of the baseline (Jain et al., 2019). Seizures were rated as convulsive if an electrographic seizure was accompanied by a behavioral convulsion (observed by video playback), defined as stages 3-5 using the Racine scale (Racine et al., 1972) where stage 3 is unilateral forelimb clonus, stage 4 is bilateral forelimb clonus with rearing, stage 5 is bilateral forelimb clonus followed by rearing and falling, and stage 6 is a stage 5 seizure with running. A seizure was defined as non-convulsive when there was electrographic evidence of a seizure but there were no stage 3-5 behaviors. If there were stage 1-2 behaviors on the Racine scale (freezing, small head bobbing) the seizure was considered non-convulsive since these behaviors were hard to discriminate from normal behavior. Records were screened manually to quantify seizures.

SeizurePro (version 2.2.14; Pinnacle Technology) or Spike2 (version 7; Cambridge Electronic Design) software was used to define characteristics of seizures. Characteristics of seizures included the type of seizure onset pattern. The majority of seizures had an onset similar to what has been described elsewhere as a low voltage fast (LVF) seizure onset in humans (Engel, 1990; Velasco et al., 2000) and in rats after systemic injection of kainic acid to induce SE (Bragin et al., 2005), after pilocarpine injection (Levesque et al., 2012), after the convulsants 4-aminopyridine or picrotoxin applied in vivo (Salami et al., 2015) or after intrahippocampal kainic acid injection in mice (Lisgaras and Scharfman, 2022). In our recordings, these seizures began with an initial sentinel spike in all leads (Figure 1, Supplemental Figure 11A1), characteristic of LVF seizures described elsewhere (Salami et al., 2015; Lisgaras and Scharfman, 2022). In many instances there was a subsequent period of low voltage, followed by large amplitude high frequency spikes. However, since it was not always low voltage, the seizure onset type was mainly characterized by a sentinel spike, a criteria also used to define LVF seizures before (Lau et al., 2022). Our LVF seizures were always accompanied by stage 3-5 convulsive behaviors.

The other type of seizure had no clear sentinel spike at its onset (Supplemental Figure 11A2). In the literature many of these seizures have an onset pattern called hypersynchronous (HYP) that is characterized by bursts of spikes at low frequency, in both humans and in rats (Bragin et al., 2005). In our mice, bursts were unclear. However, before the seizure there were bursts of fast activity (>80 Hz) that was not evident in LVF seizures (Supplemental Figure 11A1-2). Because the literature suggests that more activity at high frequencies occurs before HYP seizures compared to LVF seizures (Bragin et al., 2005; Salami et al., 2015; Avoli et al., 2016), we use the term HYP for these seizures. It should be noted that our recordings were not wideband. Wideband is important to distinguish the highest frequency oscillations (250-500 Hz; (Zijlmans et al., 2017). Therefore, we can only say our HYP seizures had higher frequency bursts before seizures relative to LVF seizures. HYP seizures were convulsive, like LVF seizures.

The third type of seizure was nonconvulsive, defined by high frequency, large amplitude activity while animals behaved normally, were frozen, or had small head movements. All seizures, including those that were nonconvulsive, occurred in each lead synchronously.

To measure the characteristics of seizures defined by the EEG, 10-15 seizures were selected starting at least 24 hours after the transition from normal water to CNO-containing water or the transition from CNO to water. Seizures were selected from days with at least 1 hour between seizures so that characteristics of one seizure were unlikely to influence the characteristics of the next seizure. Notably, even when there were many seizures/day, the interval between seizures was usually >1 hour.

Seizure duration was defined either from the EEG or the behavioral convulsion. For the **EEG seizure duration** (Figure 2), the seizure duration was measured from the sentinel spike to the end of the large amplitude, high frequency spiking that characterizes a seizure (Figure 2C). The sentinel spikes were well synchronized so any lead could be chosen to define seizure onset (Figure 2C). For some seizures, where there was less synchrony in the termination of seizures, some leads terminated up to 5 seconds after others. In these cases, the end of a seizure was defined by the electrode with the earliest termination. This electrode was selected because it typically coincided with the end of convulsive behaviors. For the **duration of the convulsive behavior**, the onset was the time when the animal started its first convulsive behavior and ended when all convulsive behavior stopped (based on video review). Also, the end of the convulsive behavior was readily defined because animals typically froze suddenly as the seizure in the EEG ended. **The latency to the convulsive behavior** was the time from the start of the seizure as defined by the EEG to the onset of convulsive behavior defined by video. It should be noted that for all seizures that had convulsive behavior the behavior always began after the EEG seizure began, i.e., after the sentinel spike.

**Figure 2.**
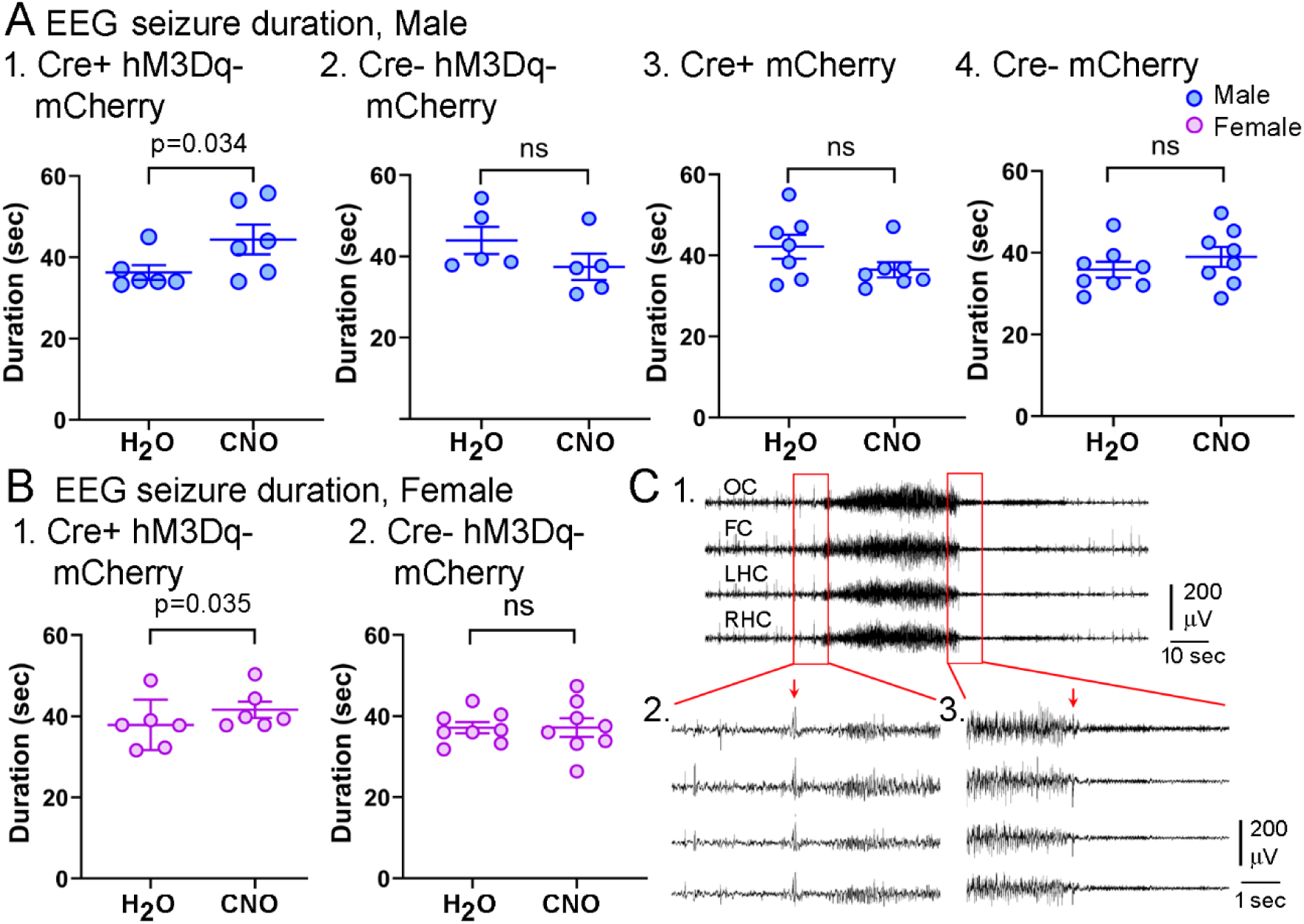
Chemogenetic activation of CA2 increases seizure duration in epileptic mice. **A.** In males, seizure duration increased when mice were provided water with CNO compared to water without CNO for Cre+ mice injected with AAV-hM3Dq-mCherry (A1, n=6) but not for controls (A2, n=5; A3, n=7; A4, n=8). Additional measurements (e.g., medians) and statistical comparisons (paired t-tests for A1-2 and A4, Wilcoxon’s test for A3) are in Supplemental Table 2. Note that for A1 the variances in the means when mice drank water without CNO vs. water with CNO were unequal. Therefore, data were log transformed to resolve the heteroscedasticity. In A1 the raw data prior to transformation are shown. The p value in A1 (p=0.034) is for the transformed data; it was also significant for data prior to transformation (p=0.041). The means, medians, maximums and 25/75 percentiles of the raw data are shown in Supplemental Table 1. The details of the paired t-test using transformed data are also shown.
**B.** Cre+ females that were injected with AAV-hM3Dq-mCherry showed a significant effect of CNO on seizure duration, although it was small (C1, n=6), but Cre- controls did not (C2, n=8). Additional measurements (e.g., medians) and statistical comparisons (paired t-tests) are in Supplemental Table 2.
**C.** Measurement of EEG seizure duration. OC, occipital cortex; FC, frontal cortex; LHC, left hippocampus; RHC, right hippocampus.

1. An example of a seizure is shown. The start and end are boxed.
2. The start of the seizure is expanded to show the sentinel spike, marking the start of the seizure (arrow).
3. The end of the seizure is expanded to show when the end was defined (arrow).

**Clusters** were defined as multiple consecutive days (≤2 days) with at least 2 seizures/day. We acknowledge that other definitions could be made and the results should be interpreted with this in mind. The **cluster peak** was assessed from the day during a cluster that had the most seizures. The number of seizures during this day (meaning the entire 24 hours) was defined as the cluster peak. The **cluster duration** was defined as the number of days between the start and end of a cluster. Thus, the first day of ≤2 seizures/day was the start and the last day of ≤2 seizures/day was the end. If the EEG recording started during a time when there were ≤2 seizures/day, no measurement of cluster duration was made because the start of the cluster was not known. If there was >1 cluster the average was used. The **number of clusters** was defined as the number of events with ≤2 seizures/day for ≤2 consecutive days. This included some clusters that were truncated by the start or end of the recording session, as long as there were >2 seizures/day for >2 consecutive days. For **intercluster interval**, the number of days between clusters was quantified, including single days when there was 1 or more seizures/day for one day. Note that when there was only one cluster, an interval could not be measured. The **number of seizure-free days** was the number of days with 0 seizures/day.

#### Acute seizures

SE was defined as the time when a convulsive seizure (≤stage 3) seizure was not followed by a return to normal behavior (grooming, eating) for several hours (Löscher, 2024). Typically animals had continuous twitching of the body but they did not have continuous convulsive seizures (≤stage 3). The duration of SE was difficult to define because it ended gradually. However, approximately 4 hours after the start of SE there was usually a relatively sudden onset of interictal spikes (IIS; Figure 4A), providing an opportunity to assess the duration of the most severe part of SE (the time until IIS began). Note that the IIS lasted overnight, as we previously described (Jain et al., 2019) and has been described by others (Mazzuferi et al., 2012; Bumanglag and Sloviter, 2018; Smith et al., 2018).

Pilot studies suggested that Cre+ mice showed more vigorous EEG activity during the onset of SE than Cre-mice. Therefore, we compared total power in Cre+ and Cre-mice using the the first 5 minutes of SE in their EEG recordings (Figure 4C). Measurement of power was done using channel 4 of the EEG, the right hippocampal channel, but results were similar for each channel. Settings were selected with 1.96 Hz bin size (FFT size, 0.512 second) and used a Hanning window. Power was measured and spectrograms were made using Spike 2 (version 7.12, Cambridge Electronic Design).

### Perfusion-fixation and histology

All methods and analyses were conducted blinded. Mice were perfused after vEEG recording. To perfuse, mice were deeply anesthetized by isoflurane inhalation followed by intraperitoneal (i.p.) injection of urethane (250 mg/kg of 250 mg/ml) dissolved in saline (0.9% sodium chloride; Cat#U2500; Millipore Sigma). After loss of a reflex to a tail pinch and loss of a righting reflex, consistent with deep anesthesia, the heart cavity was opened, and a 25 gauge needle was inserted into the heart, followed by perfusion with 10 ml saline using a peristaltic pump (Minipuls 1; Gilson) followed by 30 ml of cold (4°C) 4% paraformaldehyde (PFA; Cat# 19210, Electron Microscopy Sciences) in 0.1 M phosphate buffer (PB; pH 7.4). The brains were removed immediately and post-fixed for at least 24 hour in 4% PFA at 4°C. After post-fixation, 50 μm-thick sections were cut using a vibratome (Cat# TPI-3000, Vibratome Co.).

Sections were cut in the coronal plane and collected sequentially to be used for Cresyl violet staining to assess tissue integrity, mCherry fluorescence to detect viral expression, and immunocytochemistry with an antibody to purkinje cell protein 4 (PCP4), a marker of CA2 (Lein et al., 2005; Laeremans et al., 2013; Kohara et al., 2014; San Antonio et al., 2014) or NeuN, an antigen specific to neurons (Mullen et al., 1992; Wolf et al., 1996; Chen et al., 1997; Sarnat et al., 1998).

For Cresyl violet staining, sections were mounted on gelatin-coated (1% bovine gelatin; Cat#G9391, Millipore Sigma) slides (Cat# ZA0262; Zefon International) and allowed to dry overnight. Slides were placed in increasing concentrations of ethanol (70, 95, 100% for 2.5 minutes each) to dehydrate sections. After a second incubation in 100% ethanol for 2.5 minutes, they were rinsed in Xylene (Cat# 534056, Millipore Sigma; 2 x 2 minutes). Afterwards they were rehydrated by reversing the dehydration steps. After rinsing in ddH_2_O (3 x 5 minutes), slides were immersed in 0.25% Cresyl violet (Cat#5042; Millipore Sigma) in ddH_2_O until the desired staining intensity was reached. Then slides were placed in 4% acetic acid in ddH_2_O to reduce background staining. Afterwards slides were rinsed in ddH_2_O (3 x 5 minutes) and dehydration in ethanol was repeated. Then slides were coverslipped (Cat# 48393-106 VWR Coverglass; VWR Scientific Products Corp.) with Permount (Cat# 17986-01, Electron Microscopy Sciences). Sections were examined using an upright microscope with brightfield and fluorescence capability (Model BX51, Olympus of America). Sections were photographed using a digital camera (Model Lumera Infinity 3, Teledyne Instruments) acquired using Infinity Analyze (version 7.1.1.66, Teledyne). Figures were composed in Photoshop (version 7.0., Adobe).

For PCP4, a 2 day-long procedure was used with all steps at room temperature and using a rotator except incubation with primary antibody which was done at 4°C on a rotator. On day 1, free-floating sections were first washed in 0.1M Tris buffer (TB; 3x 5 minutes). TB contained 96.96g Tris HCL (Cat#1185-53-1; Millipore Sigma), 22.24g Tris Base (Cat#10708976001 Millipore Sigma) and 8 liters ddH_2_O. pH was adjusted to 7.6. Next, sections were washed for 10 minutes in Tris A buffer (0.25% Triton X-100, Cat#X-100, Millipore Sigma, diluted in TB) followed by 10 minutes in Tris B buffer (0.25% Triton X-100, 1% bovine serum albumin, Cat#A7906, Millipore Sigma, dissolved in TB). Subsequently, sections were incubated for 60 minutes in 10% normal donkey serum in Tris B buffer. Next, sections were incubated overnight at 4°C with rabbit anti-PCP4 antibody (1:400, Cat#HPA005792, Millipore Sigma) diluted in Tris B buffer. The antibody has been extensively validated by the Human Protein Atlas project (https://www.proteinatlas.org). It is raised to the following amino acid sequence: AGATNGKDKTSGENDGQKKVQEEFDIDMDAPETERAAVAIQSQFRKFQKKK of the human peptide and reacts to human and mouse (Gonzalez-Arnay et al., 2024). The next day, sections were washed in Tris A buffer and then Tris B buffer (10 minutes each) and then incubated in donkey anti–rabbit Alexa Fluor 488–conjugated secondary antibody (1:500; Cat#A21206, Invitrogen) diluted in Tris B. Sections were washed in TB (3 x 5 minutes). Sections were mounted onto gelatin-coated (1% bovine gelatin; Cat#G9391, Millipore Sigma) slides (Cat# ZA0262; Zefon International), and coverslipped (Cat# 48393-106 VWR Coverglass; VWR Scientific Products Corp.) with fluorescent mounting medium (Citofluor AF1; Cat#17970-25, Electron Microscopy Sciences). Slides were imaged with a confocal microscope (LSM 880, Zeiss) and Zen 3.2 software (Zeiss) to detect PCP4 and mCherry fluorescence.

For optimal visualization of double-labeled cells we used high magnification and adjusted gain and saturation so that mCherry fluorescence was not detected when using the green channel and vice-versa. Red and green colors were selected to provide a distinct color, yellow, when double-labeling occurred. We confirmed double-labeling by inspecting the red and green channels separately, ensuring that when double-labeling occurred there was fluorescence of the same location in both red and green channels. These and other aspects of our methods followed previous recommendations (Dunn et al., 2011; Elliott, 2020; Jonkman, 2020).

To quantify double-labeling we used 2 methods. We first quantified the degree that mCherry was expressed in the CA2 cell layer, which we defined by PCP4 fluorescence. Sections were used in an area that was near the injection site, approximately 1.7-2.2 mm posterior to Bregma. To this end, we cropped a region of interest (ROI) around the PCP4 fluorescence in the CA2 cell layer in ImageJ. This ROI included part of *stratum oriens* and *stratum radiatum* because the CA2 cell layer was irregular and some cells were near *stratum oriens* whereas others were near *stratum radiatum*. With the ROI, we measured the pixels that fluoresced red, green, and the total number of pixels. The ratio of red:total and green:total was calculated. Then we expressed the mean ratios for red as a percent of the ratios for green, reflecting how much mCherry fluorescence was present relative to the PCP4 fluorescence.

For the second method we set the wavelengths in the color threshold menu so that yellow pixels were measured. Afterwards we expanded the wavelengths that would be detected so that pixels of red, green, and yellow would be measured. The percent of yellow pixels of the total provided an estimate of double-labeling in the ROI.

For NeuN, free floating sections were washed in TB (3 x 5 minutes), followed by a 3 minute wash in 1% (weight/volume; w/v) H_2_O_2_ in TB. Sections were then washed in TB (3 x 5 minutes) and incubated for 60 minutes in 5% normal horse serum (Cat# S-2000, Vector Laboratories) diluted in a solution of 0.25% (volume/volume) Triton-X 100, and 1% (w/v) bovine serum albumin (#03117332001; Millipore Sigma) diluted in TB. Sections were then incubated overnight at 4°C in a mouse monoclonal antibody to NeuN (1:5000; Cat# MAB377, Millipore Sigma), diluted in a solution containing 0.25% Triton-X 100 and 1% bovine serum albumin in TB. NeuN is a well-characterized antigen (Mullen et al., 1992; Wolf et al., 1996; Chen et al., 1997; Sarnat et al., 1998) and the antibody we used has been well validated (Chen et al., 1997; Duffy et al., 2013; Corbett et al., 2017). On the following day, sections were washed in TB (3 x 5 minutes) and then incubated for 60 minutes in biotinylated horse anti-mouse IgG secondary antibody (1:500, Cat# BP-2000, Vector) diluted in a solution of 0.25% Triton-X 100, and 1% bovine serum albumin in TB. The sections were then washed in TB (3 x 5 minutes) and incubated in avidin-biotin complex for 2 hours (1:1000; Cat# PK-6100, Vector Laboratories). They were washed in TB (3 x 5 minutes) and then reacted in a solution containing 0.5 mg/ml 3, 3’-diaminobenzidine (DAB; Cat# D5905, Millipore Sigma), 40 µg/ml ammonium chloride (Cat# A4514, Millipore Sigma), 25 mg/ml D(+)-glucose (Cat# G5767, Millipore Sigma), and 3 g/ml glucose oxidase (Cat# G2133, Millipore Sigma) in TB. This method slowed the reaction time so that the reaction could be stopped when the immunoreactivity was robust but background was still low. The sections were then washed in TB (3 x 5 minutes), mounted on gelatin-coated slides and dried at room temperature overnight. The following day they were dehydrated in increasing concentrations of ethanol (90%, 10 minutes; 95% 10 minutes; 100%, 10 minutes; 100% again, 10 minutes), cleared in Xylene (10 minutes) and cover-slipped with Permount. Slides were imaged using the microscope and camera described for Cresyl violet staining.

### Statistics

Data are reported as mean ± standard error of the mean (SEM). Significance was p < 0.05. Initial sample sizes were based on prior experiments from our recent study on the effects of inhibitory DREADDs in CA2 on seizures which employed very similar methods (Whitebirch et al., 2022). During the present study several animals died or had electrodes malfunction, so the sample sizes became unequal in some cases. Particularly among females, there were animals that failed to develop SE in response to pilocarpine or failed to recover after SE. Therefore, fewer female mice were sampled in the experiments about SE so sex differences were not assessed. Power analysis was conducted *post-hoc* for all experiments. Sample sizes were based on power analysis with 1-β = 0.8 and α = 0.05 using G*Power software (Faul et al., 2009) and the values obtained for means, variances, and effect size (Cohen’s *d*). Specifically, for a comparison of main groups with significant differences, power (1-β) was >0.8 for all data with pooled sexes except for the power in the initial minutes of SE where it was 0.78. For males, power also met the criterion (0.94, seizure frequency in experimental mice, Figure 1; 0.84 for EEG seizure duration, Figure 2; 0.90 for the maximum of seizure clusters, Figure 3). For female mice, where we found significant effects the *post-hoc* power was low (0.35 for the effects of CNO on EEG sz duration, Figure 2; 0.36 for maximum cluster measurements, Figure 3) so the effects in females should be considered with this in mind.

**Figure 3.**
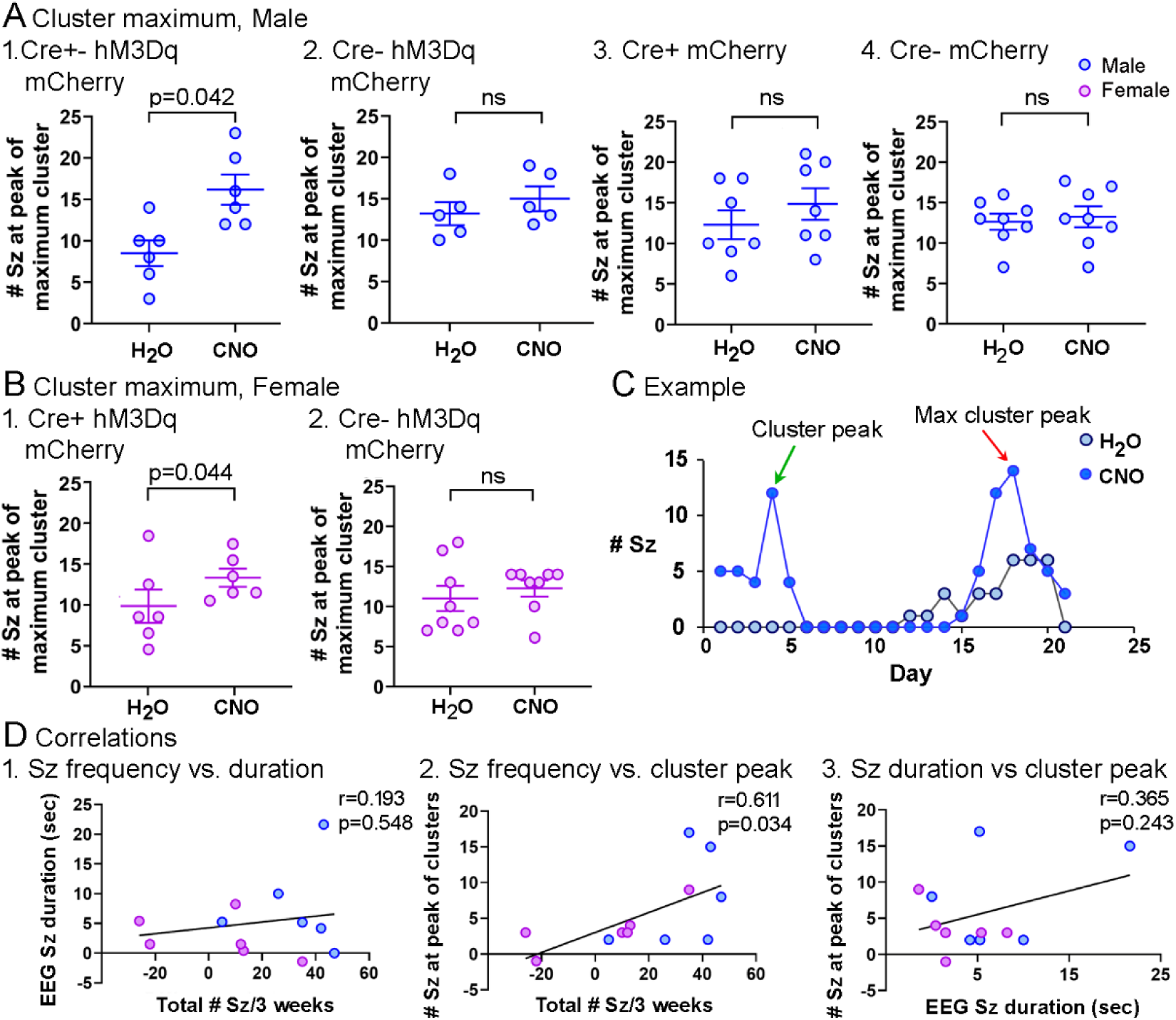
CA2 activation increases the peak of seizure clusters. **A.** The peak number for seizures/day in a cluster of seizures was determined for each cluster for each mouse and averaged to provide one value per mouse. In Cre+ males injected with AAV-hM3Dq-mCherry, the peak was greater when water was provided with CNO compared to water without CNO (paired t-test, A1, n=6). No effect of CNO was detected in controls (A2, n=5; A3, n=7; A4, n=8). Note that for A2 there are only 5 mice/group because one mouse lacked clusters (shown in Supplemental Figure 4). For A-B, all measurements and statistical comparisons are in Supplemental Table 3. **B.** In females, the peak was greater when Cre+ mice injected with AAV-hM3Dq-mCherry drank water with CNO (B1, n=6) but Cre-mice injected with AAV-hM3Dq-mCherry showed no significant effects of CNO (B2, n=8). **C.** Data from a male Cre+ mouse injected with AAV-hM3Dq-mCherry are shown to provide an example of a mouse with clusters of seizures. The number of seizures/day for each of the 21 days of a 3 week-long recording are plotted. The data for the 3 weeks on water without CNO (light blue) are superimposed on the data for the 3 weeks on water with CNO (dark blue). Arrows point to the peaks of clusters for the time when the mouse was treated with CNO. A green arrow points to the cluster with the greatest peak.

Statistical comparisons were made using Prism (version 10; Graphpad, La Jolla, CA). Data were tested for normality using the Shapiro-Wilk’s test. When there were significant departures from normality, non-parametric analyses were used. It should be noted that the Shapiro-Wilk’s test, and other tests of normality, lose accuracy when sample sizes are small. Therefore, we could have used data sets that were not normal and this is a limitation of our statistical approach. Data were also tested for homogeneity of variance using Bartlett’s test. When there was heteroscedasticity of variance, data were log-transformed. If the heteroscedasity was resolved, parametric statistics were used. If not, data were analyzed with non-parametric statistics. Parametric analyses of 2 groups used two-tailed t-tests for independent samples or a paired t-test for within-subject comparisons. For comparisons of more than 2 groups or more than one factor, one-way or multifactorial ANOVA was used. When some values were unavailable in two-way ANOVAs, a mixed effects analysis was used, and this is mentioned in the Results when it was applied. *Post-hoc* tests corrected for multiple comparisons (Tukey-Kramer). For non-parametric tests, 2 groups were analyzed by a Mann-Whitney *U* test for independent samples or a Wilcoxon signed-rank test for paired data. For correlations, parametric data were analyzed by Pearson’s r and non-parametric data by Spearman’s test.

## RESULTS

### Chronic seizure frequency is increased by chemogenetic activation of CA2 in epileptic mice

For experiments investigating chronic seizures, mice were used at approximately 8 weeks of age, when pilocarpine-induced SE leads to chronic seizures. They were injected with pilocarpine to initiate SE and virus was injected 4 weeks later, a time mice exhibit chronic seizures (Figure 1A1). By performing surgery after the vast majority of epileptogenesis had occurred, we minimized an effect of surgery on epileptogenesis. To avoid multiple surgeries, we injected virus and then, with animals still anesthetized, implanted electrodes for EEG. AAV was injected to express hM3Dq-mCherry in dorsal CA2 bilaterally. We then waited 4 weeks so there would be time for viral expression and time for recovery from implantation of electrodes. Then mice were recorded continuously with vEEG for 6 weeks. For the first 3 weeks, mice were treated either with water containing CNO or water without CNO. For the second 3 week-long period, the converse treatment was given (water without CNO for mice that had been drinking water with CNO, water with CNO for the mice that had been drinking water without CNO). After the 6 weeks of vEEG, animals were perfused for anatomical assessment of viral expression and pathology. There were some exceptions as described below.

For males, experimental mice were *Amigo2*-Cre^+/-^ (Cre+) mice injected with AAV-hSyn-DIO-hM3Dq-mCherry. Controls were *Amigo2*-Cre^-/-^ (Cre-) mice injected with AAV-hSyn-DIO-hM3Dq-mCherry, Cre+ mice injected with AAV-hSyn-DIO-mCherry, and Cre-mice injected with AAV-hSyn-DIO-mCherry. Females were *Amigo2*-Cre^+/-^ (Cre+) mice injected with AAV-hSyn-DIO-hM3Dq-mCherry. Controls were *Amigo2*-Cre^-/-^ (Cre-) mice injected with AAV-hSyn-DIO-hM3Dq-mCherry. In the text below, the name of the virus AAV-hSyn-DIO-hM3Dq-mCherry is abbreviated to AAV-hM3Dq-mCherry or referred to as eDREADD. AAV-hSyn-DIO-mCherry is abbreviated to AAV-mCherry or referred to as mCherry.

A representative example of a spontaneous seizure is shown in Figure 1B. Most seizures were robust, i.e., lasting over 15 seconds and were severe, stages 4-6. Seizure frequency of Cre+ mice injected with AAV-hM3Dq-mCherry showed a significant increase when mice had CNO in their drinking water compared to water without CNO (paired t-test, t=2.61, df=11, p=0.024; n=12 mice, 6 males, 6 females). This was not true for Cre-mice injected with AAV-hM3Dq-mCherry (paired t-test, t=1.08, df=12, p=0.300; n=13 mice, 5 males, 8 females).

The effect in Cre+ mice injected with AAV-hM3Dq-mCherry appeared to primarily be in Cre+ male mice injected with AAV-hM3Dq-mCherry rather than female Cre+ mice (paired t-tests, n=6 mice/group; Figure 1C1, 1D1) but there was higher variance in the female group, leading to low power in *post-hoc* tests (see Methods). Therefore, the results should be interpreted with this in mind.

Note that for each graph, all data for seizures during treatment with water that did not contain CNO are on the left and seizures during treatment with water containing CNO are on the right, but the temporal order of treatment was not always water without CNO followed by water with CNO (see Figure 1A1). Details of paired statistical tests as well as measurements (values for mean, SEM, median, etc.) for Figure 1 are in Supplemental Table 1. While Figure 1 displays group means, Supplemental Figure 3 illustrates how individual mice changed between treatments. Supplemental Figure 4 shows numbers of seizures each day for the entire 6 weeks of vEEG for each mouse.

The effect of CNO in male Cre+ mice injected with AAV-hM3Dq-mCherry was substantial: the mean number of seizures during the 3 weeks without CNO was 39.5 and for CNO it was 72.5, an increase of 183.5 %. In contrast, there were no significant differences for any of the male controls between the 3 weeks on water and the 3 weeks on water with CNO (paired t-tests, Cre-mice injected with AAV-hM3Dq-mCherry, n=5, Figure 1C2; Cre+ mice that were injected with AAV-mCherry, n=7, Figure 1C3; Cre-mice injected with AAV-mCherry, n=8, Figure 1C4).

In contrast to males, Cre+ females injected with AAV-hM3Dq-mCherry did not show significant effects on seizure frequency when comparing water with CNO and water without CNO (paired t-test, n=6, Figure 1D1). Cre-female controls also showed no effect of CNO (paired t-test, n=8; Figure 1D2). To shed light on the possible sex differences in effects of CNO on chronic seizures, we asked if there were sex differences in untreated mice, i.e., during the time when mice drank water without CNO. In these comparisons, Cre+ males and Cre+ females were compared, Cre-males and Cre-females were compared, and then both sexes were compared with genotypes pooled. There were no detectable sex differences in any of these comparisons in the total number of seizures during 3 weeks (Supplemental Figure 2C), suggesting no inherent sex difference could explain the sex difference in the effects of CNO on seizure frequency. There also were no detectable sex differences in EEG seizure duration when mice drank water without CNO (Supplemental Figure 5C), or sex differences in the maximum number of seizures/day during clusters (Supplemental Figure 6C). However, females drinking water without CNO had less severe convulsive behavior than males treated the same way (Supplemental Figure 8C), and shorter convulsive seizures (Supplemental Figure 9C). There was no significant sex differences in the latency to convulsive behavior (Supplemental Figure 10C). These data suggested that the convulsive behavior in untreated females was generally less severe than untreated males. Consistent with this view, several females failed to exhibit SE after pilocarpine (n=9/22, 39.1%), whereas all males did (n=26/26, 100.0%; Chi-Square test, p=0.0004). There was no effect of genotype on the number of female mice that failed to exhibit SE (Cre+, 4/10; Cre-, 5/12, Fisher’s Exact test, p>0.999). Although these data suggest a sex difference in convulsive behavior without CNO, it is not clear why this would make CNO have less of an effect on chronic seizure frequency, EEG seizure duration and the peak of clusters in females. Future studies will be required to address this issue.

In females without SE, chronic recordings were made because chronic seizures were recorded blinded to the outcome of pilocarpine administration. Interestingly, there were a small number of chronic seizures. We found no detectable effect of CNO in these females (Supplemental Figure 2A-B) consistent with the lack of detectable effects in female Cre+ mice that had SE (Figure 1D1).

Next, we asked if effects of CNO in Cre+ mice could be reversed by drinking water without CNO. To this end, a subset of mice were recorded by vEEG for a total of 9 weeks (Figure 1A2). In these animals, water without CNO was provided for the first 3 weeks, then 3 weeks of water with CNO, followed by 3 weeks of water without CNO. As shown in Figure 1E1, Cre+ mice expressing AAV-hM3Dq-mCherry showed increased seizure frequency with CNO that then decreased after mice were returned to water without CNO (RM one-way ANOVA, F(2,4)=28.00, p=0.004; n=2 males, 1 female; Figure 1E1) with a significant increase in seizures from the first 3 weeks without CNO to the 3 weeks with CNO (Tukey *post-hoc* test, p-0.008). There was a significant decline between the 3 week treatment with CNO and return to water without CNO (p=0.005), but no difference between the initial and last water treatments (p=0.787). In contrast, control mice did not differ among treatments (RM one-way ANOVA, F(2,8)=0.23, p=0.799, n=3 Cre+ males, 1 Cre-male, and 1 Cre-female; all mice were injected with AAV-mCherry; Figure 1E2). Together these data suggest that chemogenetic activation of CA2 increased seizure frequency.

### Chemogenetic activation of CA2 increases seizure duration in male and female mice

To assess the effects of CA2 activation on seizure duration, we measured the average time from the start of the electrographic seizure (EEG seizure) to its end for each mouse, in the presence of water with or without CNO. It should be noted that only a subset of all seizures were measured and these were seizures that began with a sentinel spike (Figure 1B, Supplemental Figure 11A1-2). Seizures with sentinel spikes were used because they were the vast majority of all seizures (Supplemental Figure 11A3), there was no significant differences among groups (Supplemental Figure 11A3), and because the start of the seizure was readily determined due to the sentinel spike (see Methods, Figure 1B, Supplemental Figure 11A1-2).

When sexes were pooled, Cre+ mice injected with AAV-hM3Dq-mCherry had significantly longer EEG seizure durations when they drank water containing CNO compared to water without CNO (paired t-test, t=3.48, df=11; p=0.005; n=12 mice, 6 males, 6 females). This was not true for Cre-mice injected with AAV-hM3Dq-mCherry (paired t-test, t=1.48, df=12, p=0.164; n=13 mice, 5 males, 8 females).

When sexes were separated, both Cre+ males and Cre+ females injected with AAV-hM3Dq-mCherry showed the effect of CNO in pooled mice: CNO increased seizure duration in Cre+ males (Figure 2A1) and females (Figure 2C1) injected with AAV-hM3Dq-mCherry (paired t-test, n=6 for both groups). Figure 2 shows group means and SEM, and Supplemental Figure 5 shows changes in individual mice. All measurements and statistical comparisons are in Supplemental Table 2. Control groups showed no differences in seizure duration between water with or water without CNO (male Cre-AAV-hM3Dq-mCherry, n=5; male Cre+ AAV-mCherry, n=7; male Cre-AAV-mCherry, n=8, female Cre-AAV-hM3Dq-mCherry, n=8; Figure 2A2-4, B2, Supplemental Table 2).

### Chemogenetic activation of CA2 increases clusters of seizures in epileptic mice

The seizures that were observed typically occurred in clusters, defined as ≤2 seizures/day for ≤2 consecutive days. Clusters were also found in our prior studies using the pilocarpine model (Whitebirch et al., 2022; Jain et al., 2024). It is notable that the values for the number of seizures at the peak of a cluster were similar to studies of another laboratory (Lim et al., 2018), suggesting reproducibility among laboratories. This is a clinically relevant aspect of the mouse model because seizure clusters also occur in patients with epilepsy (Bauer et al., 1992; Haut, 2006; Gidal et al., 2020; Mesraoua et al., 2021) and TLE specifically (Bauer et al., 1992).

When both sexes were pooled, the maximal number of seizures/day during a cluster increased in Cre+ mice injected with AAV-hM3Dq-mCherry when they drank water with CNO compared to water without CNO (paired t-test, t=3.47, df=11, p=0.005; n=12 mice, 6 males, 6 females). This was not true for Cre- mie injected with AAV-hM3Dq-mCherry (paired t-test, t=0.935, df=12, p=0.368; n=13 mice, 5 males, 8 females).

There also were significant differences when male and female Cre+ mice injected with AAV-hM3Dq-mCherry were analyzed separately (Figure 3A1, 3B1). CNO treatment had no effect in control mice (Figure 3A2-4, B2). Supplemental Figure 6 shows comparisons of individual mice before and during CNO. All measurements and statistics are shown in Supplemental Table 3. The duration of clusters, interval between clusters, and number of days without seizures were not significantly influenced by CNO (Supplemental Figure 7, Supplemental Table 4).

Although mean differences were significantly different in Cre+ mice injected with AAV-hM3Dq-mCherry for seizure frequency (males, Figure 1C1), EEG seizure duration (both sexes, Figure 2A1, B1), and the peak of clusters (both sexes, Figure 3A1, B1), some male and female mice showed small effects (Figures 1C1, 1D1, 2A1, 2B1, 3A1, 3B1) so we asked if mice with the greatest changes in one measurement were also those that had the greatest effects on the other measurements.

Figure 3D plots the resulting correlations. In Figure 3D1, the plot shows no significant relationship between the effect of CNO to increase seizure frequency and the effect of CNO to increase EEG seizure duration (Pearson’s r, 0.193, R^2^=0.037, p=0.548). However, there was a significant linear correlation between seizure frequency and the peak of seizure clusters (Figure 3D2, Pearson’s r, 0.611, R^2^=0.374, p=0.035). There was no correlation between EEG seizure duration and the peak of seizure clusters (Figure 3D3, Pearson’s r, 0.365, R^2^=0.133, p=0.243). These data suggest that mice could not be divided simply into those that had strong responses to CNO and weak responses. Instead, the effects of CNO appeared to depend on the measurement. The results also show that those mice with a higher total number of seizures when treated with CNO also had greater peaks to their seizure clusters, which could be explained by the contribution of higher peaks of seizure clusters to the total number of seizures.

### Characteristics of convulsive seizures

To analyze convulsive behavior, we asked if behavioral seizures were more severe on the Racine scale after eDREADD treatment. There were no significant differences in seizure severity in any of the groups (Supplemental Figure 8A-B). However, almost all seizures were severe in control conditions (stages 4-6), making it difficult to show increased severity. An interesting observation was that during treatment with water that lacked CNO, females had a lower average seizure severity compared to males, suggesting a sex difference, as mentioned above (Supplemental Figure 8C).

Measurements were also made of the duration of the convulsive behavior during a seizure, based on offline analysis of the video from vEEG recordings (see Methods).

CNO showed no effect on the duration of the convulsive behavior (Supplemental Figure 9A-B). As mentioned above, when mice were compared when only drinking water without CNO, females had shorter convulsive behavior than males (Supplemental Figure 9C).

We also measured the start of the electrographic seizure based on the EEG recording to the start of the convulsive behavior based on the video recording. We refer to this length of time as the latency to the convulsive behavior of the seizure. CNO treatment had no significant effects on this latency (Supplemental Figure 10A-B) and there were no sex differences when comparing male and female mice when they drank water without CNO (Supplemental Figure 10C). In summary, there were no detectable effects of CNO on the severity of seizures, duration of the convulsive behavior, or latency to the convulsive behavior. However, there were sex differences when mice drank water without CNO, with reduced severity and duration of convulsive behavior in females. These findings support the evidence that there are inherent sex differences in epileptic mice and provide new insight that these sex differences are primarily in convulsive behavior (see Discussion).

### Additional measurements of seizures

Other characteristics of seizures showed no effects of activation of CA2 with CNO, including the time of day of seizures, the type of seizure (defined by its onset, Supplemental Figure 11A1-3; Supplemental Table 5), the % of seizures that occurred when lights were off or on (Supplemental Figure 11B; Supplemental Table 5), and whether seizures began when mice were awake or asleep (Supplemental Figure 11C; Supplemental Table 5).

### EEG power

We also asked if EEG power during chronic seizures was influenced by CA2 activation. For this purpose, we compared LVF seizures and measured total power. Power was calculated separately for each electrode over 3 frequency bands: 0-10, 10-30, and 30-100 Hz. There were no significant differences between mice receiving water without CNO compared to water with CNO for Cre+ male and female mice injected with hM34Dq-mCherry or Cre-male and female mice injected with hM34Dq-mCherry (Supplemental Figure 12, Supplemental Table 6). It should be noted that one female Cre-mouse could not be included because of artifacts in the EEG during seizures. Therefore, the sample size for this group is 7 instead of 8 (Supplemental Figure 12, Supplemental Table 6).

### SE is altered by chemogenetic activation of CA2

The studies of chronic seizures described above prompted an interest in acute seizures. We specifically asked if activating CA2 would worsen pilocarpine-induced seizures in normal animals. To this end, Cre+ and Cre-male and female mice were injected with AAV-hM3Dq-mCherry and implanted with electrodes prior to SE, using the same methods as for mice used to study chronic seizures. After a period of 4 weeks for surgical recovery and viral expression, pilocarpine was injected to induce SE (see Methods). Control Cre-mice were injected with the same AAV. Both groups were injected with CNO 30 minutes before pilocarpine. As noted above, females often lacked SE in response to pilocarpine. The low incidence of SE in females limited the assessment of sex differences in the effects of CNO on SE.

An example of the response to pilocarpine is shown in Figure 4A-B. After the injection there was a delay and then seizures began. SE was defined as the time when a convulsive seizure (≤stage 3) was not followed by a return to normal behavior (grooming, eating) within 5 minutes. Two hours after pilocarpine injection, diazepam was injected and the amplitude of the EEG declined but did not return to normal (Figure 4A-B). Several hours later there was an abrupt transition to interictal spikes (IIS; (Figure 4A-B)) and this was used as measurement of the end of the most severe part of SE. IIS continued for the overnight hours as reported before (Iyengar et al., 2015; Jain et al., 2019).

**Figure 4.**
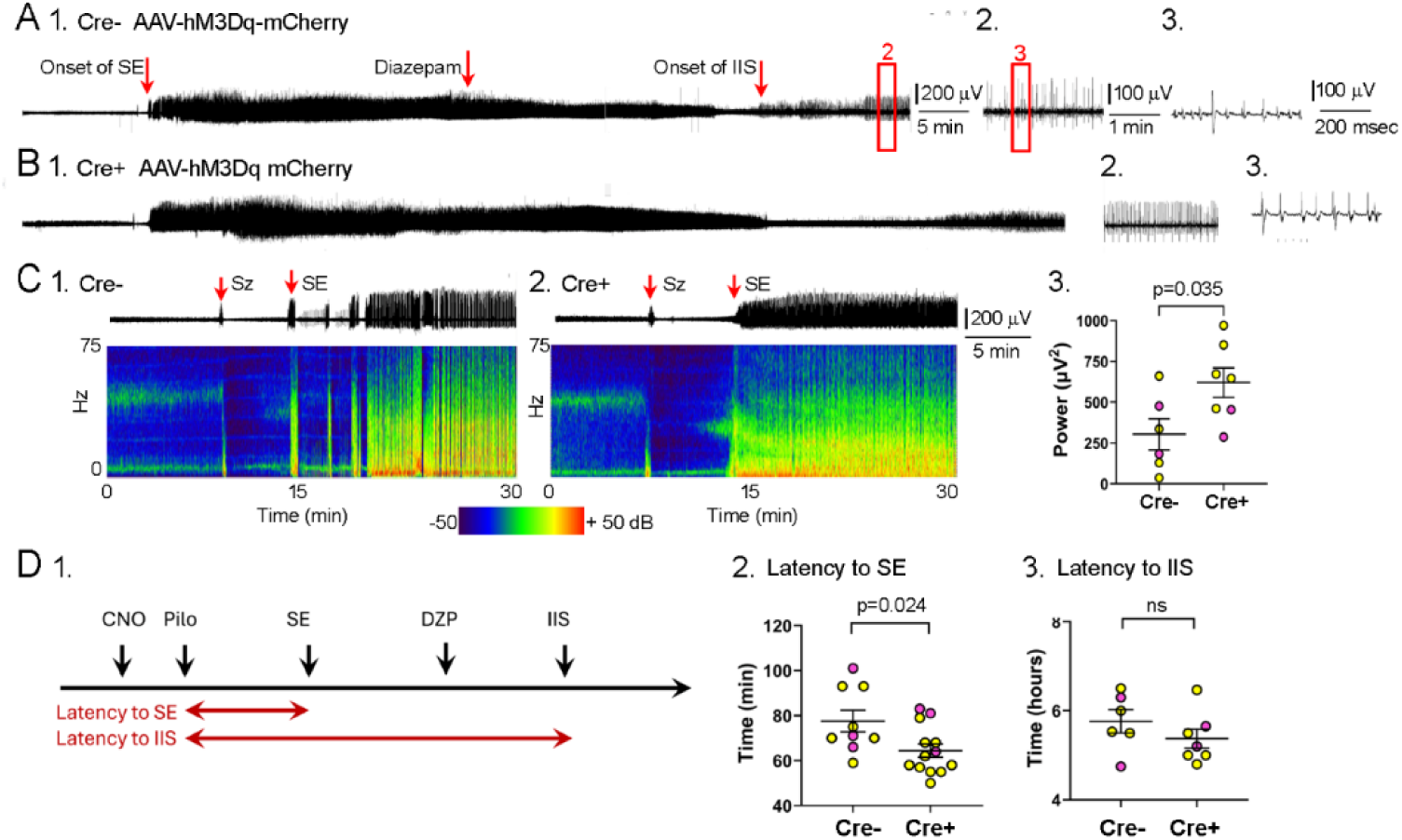
Chemogenetic excitation of CA2 shortens the latency to SE and increases EEG power during the initial phase of SE. **A.** 

1. An example of SE in a Cre+ mouse that was injected with AAV-hM3Dq-mCherry and implanted with electrodes 4 weeks before pilocarpine injection. On the day of SE induction, mice were injected with CNO (10 mg/kg, s.c.) 30 minutes before pilocarpine was injected. The EEG is shown for the channel overlying left dorsal hippocampus; the other 3 channels showed synchronous EEG activity. The onset of SE was defined by the first convulsive seizure that was not followed by a return to normal behavior within 5 minutes (red arrow). Two hours after pilocarpine injection, diazepam was injected (red arrow) and afterwards the EEG amplitude declined but did not completely return to normal. Later there was a transition to intermittent interictal spikes (IIS; red arrow). Calibration, 200 μV, 5 minutes.
2. The EEG record surrounded by the box that is labeled 2 in A1 is shown with a different time scale to make IIS more easily visualized. Calibration, 100 μV, 1 minute.
3. The box labeled 3 in A2 is shown at a different time scale to show individual IIS. Calibration, 100 μV, 200 msec.
**B.** 1-3. An example of SE in a Cre- mouse injected with AAV-hM3Dq-mCherry.
**C.** 

1. An example of SE in a Cre- mouse shown with EEG on top and the corresponding spectrogram below it. Only the time just before and after the onset of SE are shown. A seizure (Sz) occurred (first red arrow) and was followed by the onset of SE (second red arrow).
2. An example in a Cre+ mouse.
3. Total power (0-100 Hz) was calculated for the first 5 minutes of SE and showed that power was significantly increased in Cre+ mice (unpaired t-test, t=2.40, df=11; p=0.035). Sample sizes: 7 Cre+ mice (5 males, 2 females), 6 Cre-mice (4 males, 2 females).
**D.** 

1. The latency to SE and the latency to IIS are shown schematically.
2. Cre+ mice had a shorter latency to SE (unpaired t-test, t=2.44, df=20, p=0.024). Sample sizes: 13 Cre+ mice (10 male, 3 female), 9 Cre-mice (6 male, 3 female).
3. There was no significant effect of genotype on the latency to IIS (unpaired t-test, t=1.16, df=11, p=0.269). Same mice as C3.

Measurements of these events for Cre+ and Cre-mice injected with AAV-hM3Dq-mCherry are shown in Figure 4C-D. In a previous study we found that chemogenetic inhibition of mossy cells altered the power in the EEG during the initial stages of SE (Botterill et al., 2019). Therefore, we examined total power in the EEG for the first 5 minutes of SE (starting at the onset of SE in the EEG recording). Activation of CA2 in the Cre+ mice caused a significant increase in power (t=2.40, df=11, p=0.035, n= 7 Cre+, 6 Cre-; Figure 4C). We also measured the latency to SE (Figure 4D1). There was a shorter latency to the onset of SE in Cre+ mice, suggesting that activating CA2 promoted SE (unpaired t-test, t=2.44, df=20, p=0.024, n=13 Cre+, 9 Cre-; Figure 4D2). Next, we quantified the duration of SE, using the transition to IIS as the definition of the end of SE (Figure 4D1). The time between the injection of pilocarpine and the transition to IIS was not significantly affected by genotype (unpaired t-test, t=1.16, df=11, p=0.269, n=7 Cre+, 6 Cre-; Figure 4D3), suggesting the effects of activating CA2 on SE were confined to its onset. Taken together, these data are consistent with a seizure-promoting effect of CA2 activation and suggest it occurs even before the development of epilepsy.

### Targeting CA2

Next, we determined the extent of viral expression in CA2. In the first cohort we used mice that had pilocarpine-induced SE, viral injection, and were perfused 3 weeks later (Figure 5A). In a second cohort we used mice that had all procedures (pilocarpine-induced SE, viral injection, implants, 6 weeks of vEEG). They were perfused immediately after vEEG (Figure 5B). There were 5 Cre+ males injected with AAV-hM3Dq-mCherry for each cohort. In addition, we examined 13 Cre-mice after all procedures and found no detectable mCherry fluorescence (Figure 5C; 5 males injected with AAV-hM3Dq-mCherry, 6 males injected with AAV-mCherry, 2 females injected with AAV-mCherry).

**Figure 5.**
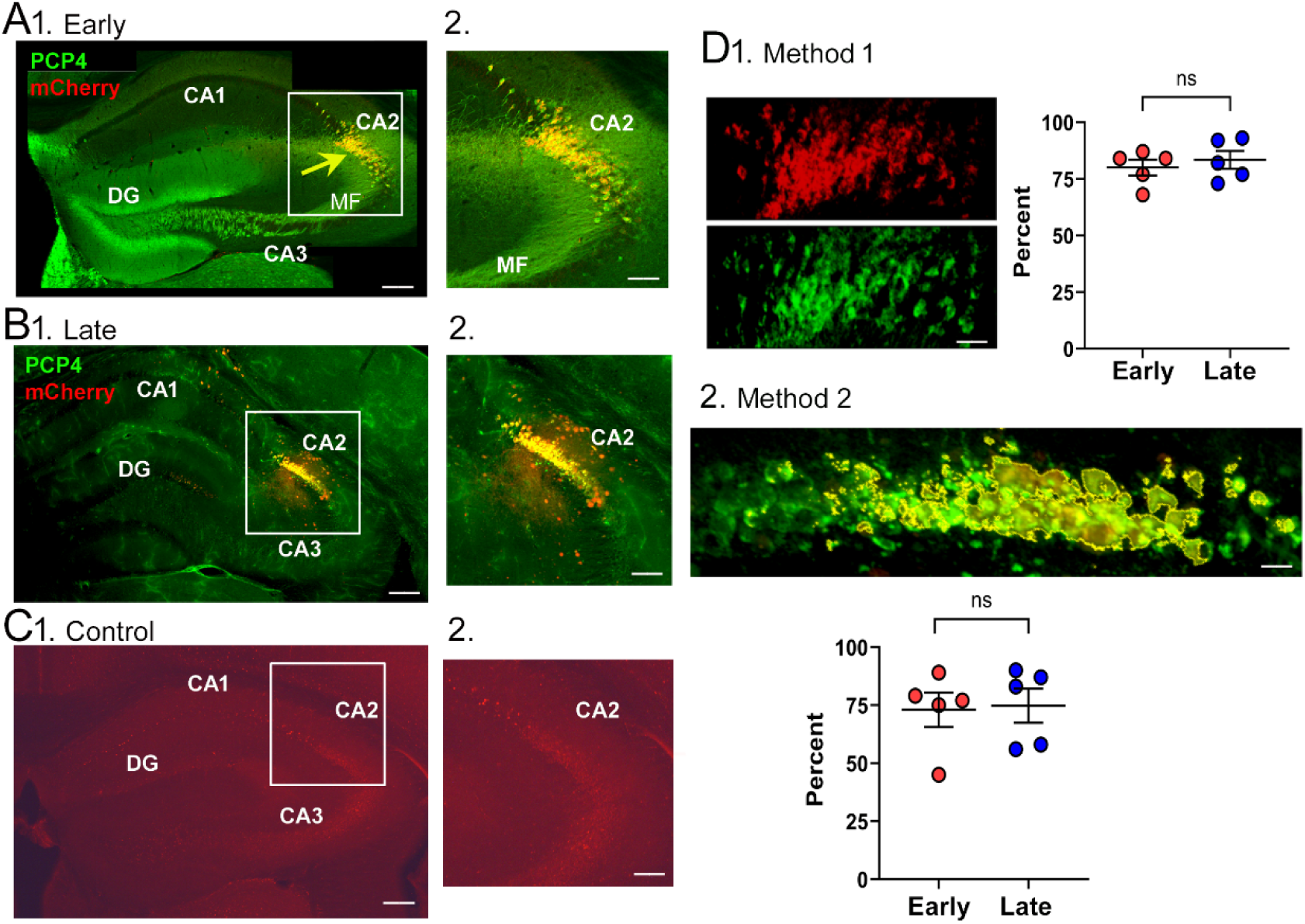
Viral expression in area CA2. **A.** Evaluation of viral expression before vEEG (early).

1. An example of a section from a male Cre+ mouse injected with AAV-hM3Dq-mCherry and stained for PCP4 (green), a marker of CA2. This mouse had pilocarpine-induced SE and viral injection and was perfused 3 weeks after viral injection. The yellow arrow points to double-labeling in CA2. Calibration, 100 µm.
2. The area in the box in A1 is expanded. Calibration, 50 µm.
**B.** Evaluation of viral expression after vEEG (late).

1. An example of a section from a male Cre+ mouse injected with AAV-hM3Dq-mCherry and stained for PCP4. This mouse had pilocarpine-induced SE, implantation of electrodes and viral injection, and was recorded 3 weeks later for 6 weeks with vEEG. Then it was perfused. Calibration, 100 µm.
2. The area in the box in B1 is expanded. Calibration, 50 µm.
**C.** Evaluation of a control mouse.

1. An example of a section from a male Cre- mouse injected with AAV-hM3Dq-mCherry. This mouse had the same procedures as the one in part B. Calibration, 100 µm.
2. The area in the box in C1 is expanded to show no mCherry expression. Calibration, 50 µm.
**D.** Quantification of viral expression using 2 methods.

1. Method 1. Pixels were measured in ImageJ in a region of interest (ROI) that was rotated and cropped from the original image. The ROI was intended to surround the CA2 cell layer and used PCP4 fluorescence to define CA2. The images that are shown are the red and green channels from the image in A. After determining the area of red pixels and green pixels and the total number of pixels in the ROI, ratios for red: total and green: total were determined. Then the mean ratios for red were expressed as a percent of the ratios for green. The high ratios (>75%) suggest that mCherry fluorescence was strong in area CA2. There was no significant difference between the ratios for early (80.0 ± 3.4%) and late (83.4 ± 4.0%) times (unpaired t-test, t=0.65, df=8; p=0.535, n=5 mice/group), suggesting stability over the time.
2. Method 2 was based on measuring the area that was yellow (double-labeled) or green (PCP4) in ImageJ. The first procedures to rotate and crop the image were the same as Method 1. The image, from B1, shows the threshold for pixels that were considered both red and green (yellow outlines). After determining the area of yellow pixels relative to the pixels in all wavelengths, the ratio was expressed as a percent and is plotted for the same mice as in D1. There was no significant difference between the early (73.0 ± 7.4%) and late (74.8 ± 7.4%) times (unpaired t-test, t=0.17, df=8; p=0.867, n=5 mice/group), again suggesting stability. Thus, both methods suggest strong, stable viral expression in CA2.

In the Cre+ mice (Figures 5A-B), we quantified double-labeling for mCherry and the CA2 marker PCP4 in 2 ways (see Methods). First, the degree of overlap between mCherry and PCP4 fluorescence was determined. The average ratio of mCherry/PCP4 was >75% percent (Figure 5D1), suggesting that experimental mice had strong expression of mCherry in CA2, although it was not 100%.

Next, we quantified double-labeling by comparing all pixels that were yellow (double-labeled) relative to the pixels that fluoresced red (mCherry), green (PCP4), and yellow. To define CA2 we again cropped an area around PCP4 fluorescence. The area that was yellow, expressed as a percent of the area measured for all wavelengths, was >75% (Figure 5D2). The results suggest strong double-labeling in CA2 in the Cre+ experimental mice but, again, it was not 100%.

We also compared the early and late timepoints and found no significant differences (Figure 5D1-2), suggesting that there was stable expression throughout the procedures. This stability is important because if expression waned with time after injection, the results would be difficult to interpret.

### Neuronal loss in epileptic mice

Previous studies have shown that CA2 is relatively resistant to pilocarpine-induced SE (Whitebirch et al., 2022). In contrast, a characteristic pattern of neuronal loss occurs in the hilus, CA3, and CA1 (Jain et al., 2019; Whitebirch et al., 2022, 2023). To confirm that CA2 neurons survived SE and chronic seizures and to characterize neuronal loss in hippocampus in our experiments, mice were perfusion-fixed after 6 weeks of recording. Using Cresyl violet-stained sections, we measured the area of the dorsal hippocampus from individual sections, selecting 3 that were approximately 300 μm apart, as shown in Figure 6A-B. In contrast to saline controls (Figure 6A), pilocarpine-treated mice had significant damage, particularly in area CA3 (arrows, Figure 6B). Comparisons of the area of the hippocampus (Figure 6C1) showed that the most rostral section, section 1, was significantly larger in saline controls than pilocarpine-treated mice. The results suggested significant loss of area in pilocarpine-treated mice for the most rostral hippocampus. Next, we confirmed that there was no significant difference in this effect for Cre+ and Cre-mice (Figure 6C2). Figure 6D illustrates the methods to define hippocampal area. Because hilar cell loss was common to all mice, the neurons in the hilus were quantified in sections from dorsal hippocampus at approximately 1.6-1.8 mm posterior to Bregma. As shown in Figure 6E, only the large cells with the morphology of neurons (exemplified by the cell marked by the arrow in 6E) were counted and small cells with the morphology of glia were not. For comparison, naïve Cre-mice that had sterile saline instead of pilocarpine were quantified. The results showed that the number of hilar cells was reduced in all pilocarpine-treated mice compared to controls, there were no significant differences between Cre+ and Cre-mice, and no sex differences (Figure 6F). The results suggest that the effects of CNO described in Figures 1-4 were not biased by differences in the degree of neuronal damage between groups.

**Figure 6.**
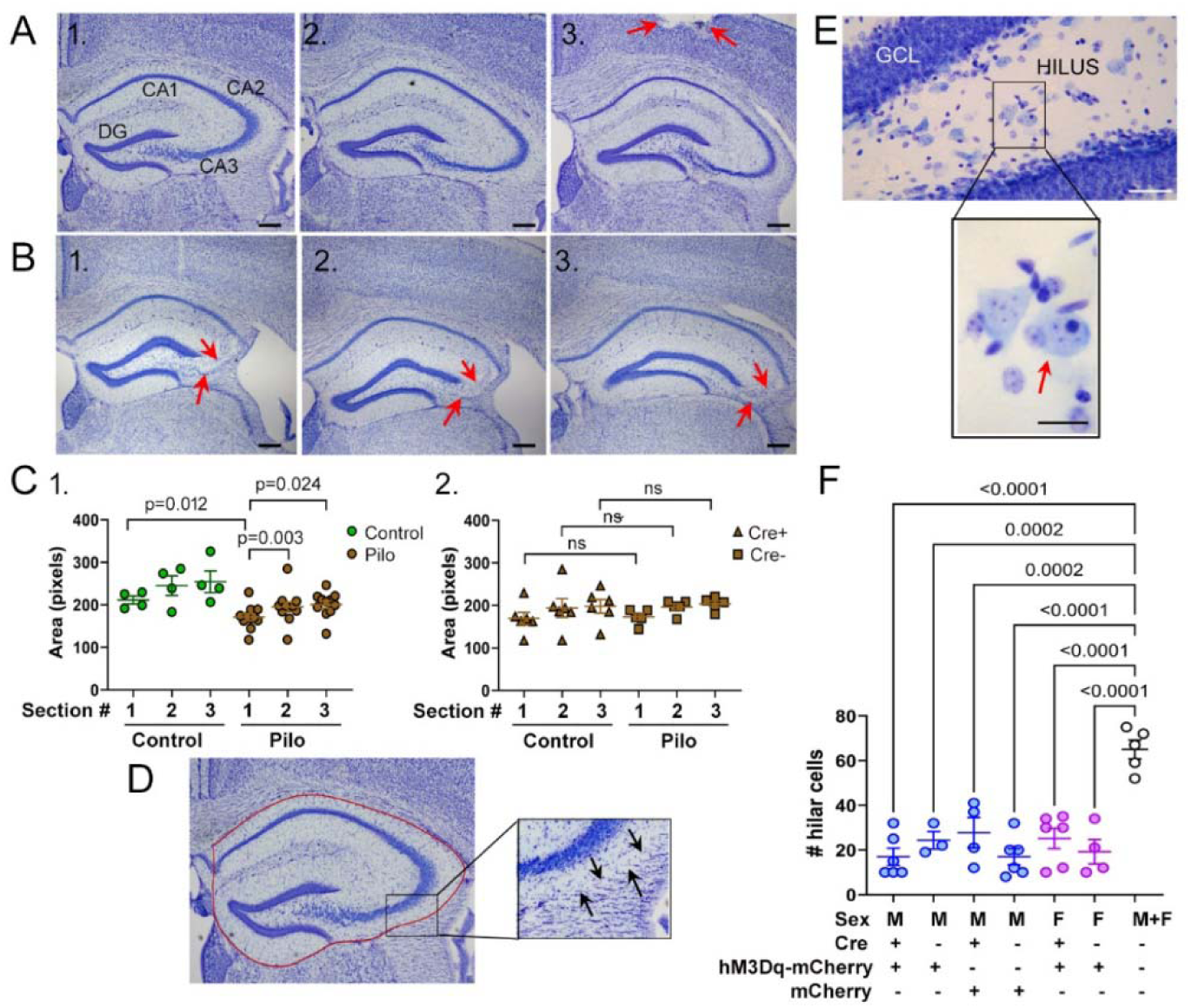
Hippocampal area decreased after SE. **A.** Sequential coronal sections from a control mouse treated with saline instead of pilocarpine. Sections were stained with Cresyl violet and are shown from the most septal (1), intermediate (2), to the most caudal (3) section. Sections were approximately 300 μm apart. Mice were implanted with electrodes and perfused at the same time as mice that had pilocarpine (14 weeks after saline; Figure 1A1). Arrows marks damage near the site of one of the EEG electrodes. DG, dentate gyrus. Calibrations, 100 μm.
**B.** Sections from a mouse that was treated with pilocarpine and had SE. Sections were selected from the same locations along the septotemporal axis as those in A. There is substantial loss of the CA3 pyramidal cells (arrows). Calibrations, 100 μm.
**C.** Comparison of the area of the hippocampus measured by tracing its borders (shown in D). Saline-treated control mice (Con) are compared to pilocarpine-treated mice (Pilo).

1. A two-way ANOVA with treatment (Con, Pilo) and section number (1-3) as main factors was significant (treatment, F(1,13)=8.23, p=0.013; section number, with Geisser-Greenhouse correction, F(1.48, 19.35)=7.74, p=0.006). For the first section, there was significantly less hippocampal area in pilocarpine-treated mice (*post-hoc* Tukey-Kramer tests, p=0.012). Also, in pilocarpine-treated mice, the first section had smaller area than the other sections (1 vs. 2, p=0.003; 1 vs. 3, p=0.024). Sample size: saline, 4 mice; pilocarpine, 11 mice.
2. Comparison of Cre+ and Cre- pilocarpine-treated mice by two-way ANOVA did not show significant differences in genotype (F(1,9)=0.037, p=0.851). *Post-hoc* tests showed no differences between genotypes for section 1, 2, or 3 (Tukey-Kramer tests, all p>0.05). Sample size: Cre+, 6 mice; Cre-, 5 mice.
**D.** Methods for quantification of hippocampal area. The same section as A1 is shown with a red line illustrating the borders of the hippocampus. The inset shows the ability to discriminate the borders of the hippocampus and extrahippocampal areas by examining the distribution of Cresyl violet-stained cells at high power, where the distribution was distinct (arrows point to the border of the alveus with the fimbria).
**E.** A high magnification of the hilar region shows large cells corresponding to neurons and small cells reflecting other cell types. The inset shows the large cells, exemplified by the one marked by the arrow, that were quantified to estimate the number of surviving hilar neurons.
**F.** The mean number of hilar neurons was averaged for sections 1-3 in each mouse. Comparisons were made between saline-treated controls (Con, white symbols, last column) and pilocarpine-treated epileptic mice (all other columns). There were significantly fewer hilar neurons in all epileptic mice compared to controls (one-way ANOVA (F(6,27)=13.83, p<0.0001). Because we did not detect significant differences between males (blue symbols) and females (pink symbols), sexes were pooled for the controls.

To assess neuronal loss, we analyzed the pyramidal cell layers using NeuN immunocytochemistry. This assessment was made based on the observations of gaps in the pyramidal cell layers, reflecting a loss of neurons (Figure 7A2-3). Therefore, the total length of the cell layers was measured, subtracting the gaps. Landmarks discriminated the borders of CA1, CA2, and CA3 (Figure 7C). The results showed significantly smaller pyramidal cell layer length in CA1 and CA3 but not CA2. The results are consistent with substantial loss of neurons in hippocampus with relative resistance of CA2.

**Figure 7.**
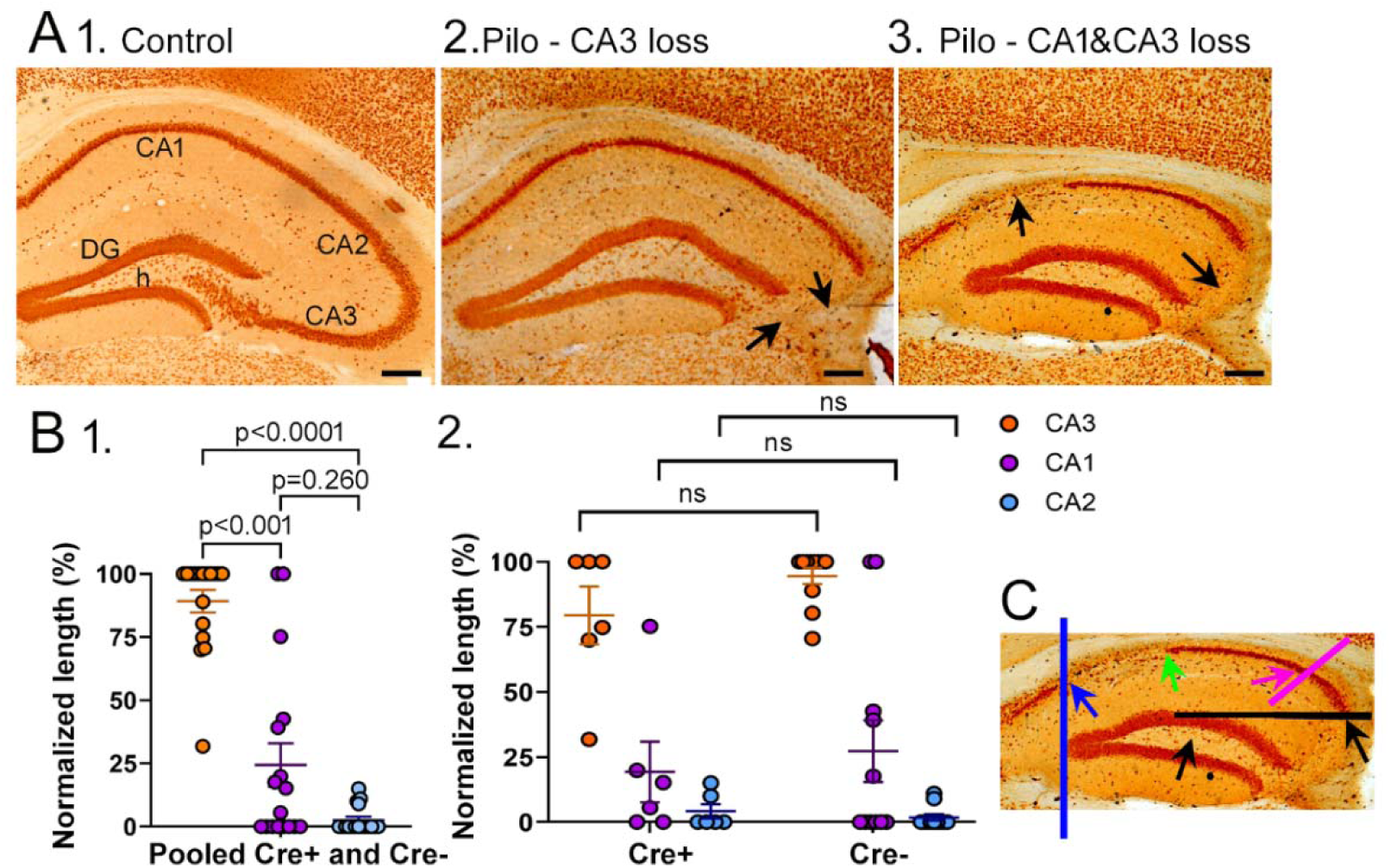
Hippocampal neuronal loss after SE. **A.** Sections from saline (Control) or pilocarpine (Pilo)-treated mice stained with an antibody to the neuronal antigen NeuN. DG, dentate gyrus; h, hilus. Mice underwent procedures shown in Figure 1A1 and were perfused at the same times, 14 weeks after saline or Pilo. Calibrations, 100 μm.

1. A section from a saline control mouse.
2. A section from a mouse that had pilocarpine-induced SE. Neuronal loss in area CA3 is marked by arrows.
3. A section from another mouse that had SE shows neuronal loss in both area CA1 and CA3 (arrows). The pattern of cell loss, which includes resistance of area CA2 and the DG granule cells to neuronal loss, is described as hippocampal sclerosis (Margerison and Corsellis, 1966; Falconer, 1974; Babb et al., 1984; Scharfman and Pedley, 2007; Blümcke et al., 2012).
**B.** Quantification of neuronal loss in the hippocampal cell layers. The layers were traced in ImageJ and then the length that lacked detectable NeuN staining was expressed as a percent of the total length. All mice had SE, AAV injection and electrode implantation, and were recorded by vEEG for 6 weeks before perfusion-fixation as shown in Figure 1A1.

1. For both Cre+ and Cre-mice (data pooled), there was more neuronal loss in area CA3 than CA1 and very little neuronal loss in area CA2, consistent with its relative resistance. A Friedman’s test showed significant differences in subfields (p<0.0001; Friedman’s statistic, 29.93). Dunn’s multiple comparisons tests showed CA3 had more damage than CA1 (p=0.003) and CA2 (p<0.0001). Sample sizes: 6 Cre+ mice (3 AAV-hM3Dq-mCherry, 3 AAV-mCherry), 11 Cre-mice (2 AAV-hM3Dq-mCherry, 9 AAV-mCherry). Some mice were not evaluated because of technical problems (damage while perfusing, removing the implant or during sectioning).
2. Comparisons of Cre+ and Cre-mice showed no significant effect of genotype (CA3: Mann-Whitney *U* test, p=0.211, *U*=22; CA1: Mann-Whitney *U* test, p=0.952, *U*=32; CA2: Mann Whitney *U* test, p=0.3529, *U*=27). Same mice as B1.
**C.** Methods for quantification. The methods used to define the cell layers and measurements to quantify neuronal loss are shown. For CA1, the length of CA1 that had no detectable NeuN was measured between the blue and green arrows. The total length of CA1 was measured between the blue and pink arrows. The length with neuronal loss was expressed as a percent of the total length. Other subfields were measured in an analogous manner. The borders of the subfields were defined as follows: the border of CA1 and subiculum was defined by a vertical line through the apex of the DG granule cell layer (blue line). The end of CA1 and the start of CA2 was defined by the point (pink arrow, pink line) where pyramidal cell size and width of the layer changed, with CA2 pyramidal cells larger than CA1 (Dudek et al., 2016). The border of CA2 and CA3 was defined by a line in the medio-lateral plane starting at the most dorsal point of the DG granule cell layer and ending at the alveus (black line). This definition was chosen because the more common definition, i.e. the transition from CA2 to CA3 pyramidal cell morphologies, was not possible when the part of CA3 adjacent to CA2 was lost. The end of CA3 in the hilus was defined as the midpoint between the lateral tip of the superior blade and the apex of the DG (black arrow in the hilus).

## DISCUSSION

### Chronic seizure frequency is increased by chemogenetic activation of CA2 in epileptic mice

In Cre+ mice injected with AAV-hM3Dq-mCherry, there was a significant increase in seizure frequency when CA2 was activated by CNO compared to periods without CNO and to results in control groups with or without CNO. These data strongly suggest that chemogenetic activation of area CA2 increases seizure frequency. The results are supported by the use of several control groups. Moreover, in a subset of the Cre+ mice injected with AAV-hM3Dq-mCherry the initial seizure frequency with water alone was significantly increased when CNO was added to the drinking water and was reversed when water without CNO was resumed. When sexes were separated, Cre+ males injected with AAV-hM3Dq-mCherry showed an effect, but Cre+ females injected with AAV-hM3Dq-mCherry did not. However, the variance in females led to a low power for that group so a definitive sex difference is unclear.

### EEG seizure duration

CNO also increased seizure duration in Cre+ males and females injected with AAV-hM3Dq-mCherry. This finding is interesting because seizure duration is likely to reflect the ability of seizures to reverberate or repeatedly pass from cortex to hippocampus and back to cortex. On the other hand, seizure frequency is likely to reflect the site where seizures are generated. Thus, CA2 may be involved in seizure initiation and may also facilitate reverberation of seizures.

A role in seizure initiation may be related to the normal ability of CA2 to synchronize neural activity in downstream CA1, based on the ability of CA2 to produce sharp waves ripples in CA1 (Oliva et al., 2016; Piskorowski and Chevaleyre, 2018; Robert et al., 2018). Although CA2 also has strong GABAergic inhibition (Mercer et al., 2012), after pilocarpine-induced SE inhibition is decreased in CA2 (Whitebirch et al., 2022), which could lead to more synchronized excitatory activity and ultimately seizures.

A role of CA2 in reverberatory activity is suggested by what is known about its circuitry. CA2 normally activates contralateral CA2 and ventral hippocampus (Meira et al., 2018). After pilocarpine-induced SE, there is an increase in granule cell input to CA2, an increase in CA2 output to CA1, and as mentioned above, GABAergic inhibition declines (Whitebirch et al., 2022). These changes would place CA2 in a position to promote seizure activity that passes from cortex through the hippocampal circuitry and ultimately back to cortex.

### Clusters of seizures

The activation of CA2 with CNO increased the maximum number of seizures/day during a cluster of seizures (the “peak” of a cluster). This is a significant result that is relevant clinically because seizures clusters in humans can be debilitating. However, because it is largely unknown why seizures cluster it is hard to suggest how CA2 would exacerbate the number of seizures during a cluster. One possibility is suggested by the hypothesis that epileptiform events lead to short-term plasticity which promotes further seizures (Ergina et al., 2021; Yang et al., 2021; Marin-Castaneda et al., 2024), an example of maladaptive plasticity. One might think CA2 would not be relevant to a possible mechanism for clustering that requires synaptic plasticity, because normally area CA2 has a limited degree of long-term potentiation or LTP (Zhao et al., 2007; Lee et al., 2010). However, degradation of perineuronal nets rescues the CA2 deficit in LTP in a mouse model of Rett syndrome (Carstens et al., 2016; Carstens et al., 2021). In addition, behavioral stress rescued short-term LTP in CA2 (Carstens et al., 2021).

These studies are relevant because, in epileptic mice, perineuronal nets in CA2 are reduced (Whitebirch et al., 2023), and behavioral stress is increased (Mazarati et al., 2009; Hooper et al., 2018; Salpekar et al., 2020). Therefore, synaptic plasticity in CA2 of epileptic mice may be more robust than normal and foster maladaptive plasticity like seizure clusters. If this were the case it would explain why eDREADD activation of CA2 would influence clusters.

### Relationships between seizure frequency, EEG seizure duration, and the peak of clusters

There was a significant correlation between seizure frequency and the peak of clusters. Thus, when comparisons were made between Cre+ mice injected with AAV-hM3Dq-mCherry that drank water with vs. without CNO, mice that had a relatively large increase in seizure frequency while drinking water with CNO also had a relatively large increase in the peak of clusters. One explanation is that the number of seizures at the peak of clusters is likely to influence the total number of seizures in the 3 week period, with larger peaks contributing to a greater total number of seizures.

There was no significant correlation between seizure frequency and EEG seizure duration or EEG seizure duration and the peak of clusters. These results suggest the effects of CA2 activation on seizure frequency and cluster peaks may be mediated by a mechanism distinct from that which prolongs seizure duration. Indeed, seizure frequency and seizure duration may be anticorrelated because an increase in seizure duration may decrease seizure frequency due to the longer time needed to recover from a seizure before another is initiated. A prolonged seizure duration may also limit the number of seizures that can be generated during a cluster for the same reason. Exactly how long a seizure needs to be to have such an inhibitory effect is not known and may not be linear, making this possibility speculative at the present time.

The result that only one correlation was statistically significant is also valuable, although it should be treated cautiously because all trendlines showed a similar relationship even though only one was statistically significant. Thus, the trendlines suggested an increase in one measure was associated with an increase in the other measure. Nevertheless, because only one was statistically significant, it is worth considering the implications of these statistical findings. One implication is that the data were not binary, i.e., there was no sign of a few animals responding well in all ways and another group that did not response well to any measure. That is important because if it had occurred it suggests a possible technical confound preventing a subset of mice from responding well. Instead, the findings suggest a dissociation between effects of CNO on different aspects of chronic seizures.

### Severity of seizures

To analyze convulsive behavior we asked if seizures were more severe on the Racine scale during eDREADD activation. There were no significant differences in seizure severity, but almost all seizures were severe in control conditions so it would be difficult to show increased severity. Measurements were also made of the latency from the start of the EEG seizure to the start of the convulsive behavior. CNO showed no significant effects.

A number of other characteristics of seizures were not altered by CA2 activation, such as the time of day of seizures. Thus, the percent of seizures that occurred when lights were off or on was not significantly influenced by CA2 activation. Whether seizures began when mice were awake or asleep was not significantly affected either. These data are consistent with the lack of direct connections between CA2 and areas that control the circadian rhythm (Cui et al., 2013; Dudek et al., 2016).

### EEG power

To determine whether EEG power during seizures was influenced by chemogenetic activation of CA2 we compared LVF seizures since they were the predominant type of seizures, and quantified power for the low (0-10 Hz, including delta, theta), 10-30 Hz, including alpha, beta) and high frequency bands (30-100 Hz, gamma). These calculations were made for each lead to ask if the hippocampal or cortical power might be different, but there were no significant differences. The results are consistent with the lack of effect of CNO on seizure severity, since power could be considered a proxy for seizure severity. The fact that power between 30-100 Hz was not affected is interesting because CA2 has been suggested to influence gamma oscillations in hippocampus of normal mice (Alexander et al., 2018). The fact it did not in the epileptic mice may be due to the loss of GABAergic neurons in these mice (Whitebirch et al., 2023) and supports the idea that CA2 properties are altered in the epileptic brain relative to normal conditions (Tulke et al., 2019; Freiman et al., 2021; Whitebirch et al., 2022; Kilias et al., 2023).

### Role of CA2 in promoting SE

The studies of chronic seizures lead to questions about acute seizures, namely, does activation of CA2 promote an acute seizure in the normal brain? It might because CA2 appears to induce synchronization of subsets of CA3 pyramidal cells, leading to sharp wave-ripples (Oliva et al., 2016). To address this question directly, we compared Cre+ and Cre-mice injected with AAV-hM3Dq-mCherry during pilocarpine-induced SE. We found that the latency to SE was shorter and initial power was greater in mice pre-treated with CNO, suggesting that CA2 activation does promote SE. Therefore, CA2 activation facilitates seizures in both normal and epileptic conditions.

### Role of CA2 in epilepsy

It is remarkable that activation of a small area like dorsal CA2 can have significant effects on seizures. How would such a small area exert robust control of seizures? Studies of the projections of CA2 neurons suggest a reason. CA2 projects bilaterally to CA3 and to CA1. That widespread influence is likely to contribute to its robust regulation of seizures. Moreover, there is increased excitatory synaptic input relative to synaptic inhibition in CA2 in the pilocarpine model (Whitebirch et al., 2022). One of the enhanced excitatory inputs was from the mossy fiber projection from the granule cells, which is consistent with findings in the intrahippocampal kainic acid (IHKA) model and patients with intractable TLE that there is increased mossy fiber sprouting into CA2 (Freiman et al., 2021; Kilias et al., 2023). Reduced inhibition appears to be due to a loss of GABAergic neurons in CA2, which is primarily due to loss of cholecystokinin-expressing GABAergic neurons in the pilocarpine model (Whitebirch et al., 2023). In the IHKA model, reduced GABAergic neurons have also been reported (Kilias et al., 2023). In addition, theta oscillations are reduced (Kilias et al., 2023), which is relevant because theta oscillations have been suggested to inhibit seizure activity in rodents with epilepsy (Miller et al., 1994).

Another potential contributing factor is an increase in the neurotrophin brain-derived neurotrophic factor (BDNF) in CA2 neurons in the IHKA model of TLE (Tulke et al., 2019). BDNF can promote glutamate release, decrease GABAergic inhibition, and LTP in area CA1 and the dentate gyrus (Kang and Schuman, 1995; Scharfman, 1997; Frerking et al., 1998; Messaoudi et al., 1998; Asztely et al., 2000; Lu et al., 2014). It is also proconvulsant in epilepsy (McNamara and Scharfman, 2012). These mechanisms may contribute to the role of CA2 to promote seizures.

Recurrent collaterals and connections between CA2 PCs of each hemisphere may also play a role (Meira et al., 2018; Okamoto and Ikegaya, 2019). Thus, activating a subset of CA2 neurons may lead to many more CA2 neurons that are activated by recurrent excitation. Moreover, CA2 projects to contralateral CA2, which has been confirmed in the IHKA model (Kilias et al., 2023). In that model, it was shown that coherence of CA2 between the two hemispheres increased as epilepsy developed and it was suggested that this increase in coherence may contribute to epileptogenesis (Kilias et al., 2023). Therefore, by expressing eDREADDs bilaterally in CA2 we may have promoted positive feedback pathways between CA2 regions locally and between the two hemispheres, promoting synchronization and therefore seizures.

### Neuronal loss in epileptic mice

Our assessments of hippocampal neuronal loss after SE showed a significant reduction in hippocampal area, indicative of shrinkage. Of the subfields, area CA1, CA3 and the hilus showed significant neuronal loss. In contrast, CA2 was relatively resistant, consistent with past studies showing that CA2 is spared in animal models of TLE and in humans with TLE. (Babb et al., 1984; Leranth and Ribak, 1991; Sloviter et al., 1991; Whitebirch et al., 2022, 2023; Kilias et al., 2023). It remains unclear why CA2 is resistant. Initial hypotheses focused on the mechanisms that limit calcium entry into neurons during excitotoxicity, since epileptogenesis is typically initiated by an event that triggers excitotoxicity such as SE (Scharfman and Schwartzkroin, 1989; Sloviter, 1989; Fairless et al., 2019)). One example is the high expression in CA2 of calbindin D28K since it is a calcium binding protein that theoretically can bind excess intracellular calcium (Leranth and Ribak, 1991). However, other hypotheses have been suggested based on criticisms of the original calcium binding protein hypothesis (Mockel and Fischer, 1994; Klapstein et al., 1998). Thus, resistance of CA2 has been attributed to expression of proteins in CA2 that inhibit neuronal activity such as *Amigo2* (Laeremans et al., 2013). CA2 neurons also express unique potassium channels, have strong GABAergic inhibition, and extensive perineuronal nets (Mercer et al., 2012; Dudek et al., 2016; Donegan et al., 2020; Whitebirch et al., 2023). These characteristics could lead to reduced excitation and therefore reduced excitotoxicity during brain injury.

### Sex differences

We found several sex differences in both the animals that had no CNO and in the effects of CA2 manipulations. Regarding sex differences in the absence of CNO, we found that females did not have SE as often as males, despite receiving the same dose of pilocarpine as males. In the past we also found that pilocarpine was less effective in eliciting SE in female than male mice (Jain et al., 2024), as found by others (Buckmaster and Haney, 2012; Hong et al., 2021; Wick et al., 2025). However, it is notable that many studies do not report sex differences in SE. This difference may reflect differences in methodology, including the fact that other studies used a higher dose of pilocarpine, although our past studies did not reveal a dose-dependent increase in seizures in females. Another explanation for the lack of sex differences in some studies is that females may not have had housing that fosters estrous cycling, leading to altered hormone levels that in turn influence excitability and epilepsy, as discussed elsewhere (Scharfman and MacLusky, 2006, 2014a). Nevertheless, it is important to note that the literature is varied in its evidence for sex differences in pilocarpine-induced SE, and more research will be needed to understand the variability.

Remarkably, despite a lack of SE, some female mice developed epilepsy. This finding is surprising because past studies suggest that in males, SE is required for epileptogenesis (Brandt et al., 2015; Chen et al., 2017). However, one study of rats did find that males without SE developed chronic seizures, although that occurred after a long delay (8 months; Navarro Mora et al., 2009). Therefore, it appears that females without SE develop chronic seizures earlier than males, but both males and females can develop chronic seizures without SE.

Why females without SE develop chronic seizures more readily than males is unknown. One possible explanation is that less pilocarpine-induced acute seizure activity may be required in females than males for epileptogenesis. This idea is supported by studies showing that some characteristics of the hippocampus that support activity-dependent plasticity of neural circuits are greater in females. For example, female rodents have greater BDNF levels than males (Harte-Hargrove et al., 2013; Lu et al., 2014; Scharfman and MacLusky, 2014b). BDNF is especially relevant to this study because, as mentioned above, it promotes seizures (MacNamara and Scharfman, 2012).

After SE, females and males showed differences in their chronic seizures. There were differences in the durations of clusters, severity of convulsions and the duration of convulsive behavior was reduced in females relative to males. This is interesting because it supports the idea that BDNF, and estrogen, are proconvulsant but also protective (MacNamara and Scharfman, 2012). Another possibility is related to the predisposition of females for spreading depolarization (Brennan et al., 2007; Kudo et al., 2023; Boyce et al., 2025), which often occurs at the end of convulsive seizures and is coincident with postictal depression (Ssentongo et al., 2017; Tamim et al., 2021; Jain et al., 2024). Interestingly, we showed previously that postictal depression was more severe in females (Jain et al., 2024). Therefore, females may have a greater susceptibility to spreading depolarization and postictal depression. The result might lead to shorter convulsions that do not continue to reverberate and attain greater severity. Alternatively, different levels of hormones in females and males may lead to differences in excitability of those brain areas that control convulsions (Giorgi et al., 2014).

This study also found that CNO affected seizure frequency in males but there was no detectable effect in females. This result has to be treated cautiously because the female data showed high variability. However, if it reflects a sex difference in the effects of CNO it is intriguing. The implication is that sex differences exist in area CA2 that will influence epilepsy, although to our knowledge only one study regarding sex differences in CA2 has been published and was not about seizures (Jabra et al., 2024). Therefore, more experiments will be required to understand the sex differences in our results.

### Additional considerations

Despite the overall effects of CA2 activation on seizure frequency, duration and the maximum seizure frequency within a cluster, some animals showed weak effects. In addition, control mice with normal CA2 function showed variability, making seizure characteristics in some experimental mice overlap with the values for controls. Therefore, the results should be interpreted with these reservations in mind. However, when one uses paired comparison of animals treated with water alone versus the same mice treated with CNO, the effects become clear. The strength of the paired comparisons is likely to be due to the variability within the pilocarpine model and in the sexes.

Although there were effects of CNO on seizure frequency, seizure duration and the peak of seizure clusters, several other measurements of seizures were not affected. These findings can potentially be explained by the differences in mechanisms underlying each aspect of chronic seizures. For example, seizure frequency is influenced by mechanisms that initiate a seizure, which in the case of limbic seizures involve hippocampal subfields (Gale, 1992), an idea that has also been suggested for the pilocarpine model (Jefferys, 2014). However, for seizure severity, at least the differences between stages 4, 5, and 6 seizures that we measured, severity depends more on spread to extratemporal areas including the brainstem. Indeed the transition to running (stage 6) has been suggested to be dependent on midbrain and brainstem structures (Gale, 1992; Walker et al., 1999; Faingold, 2012).Thus, if seizure initiation is influenced by area CA2, CNO would be expected to influence seizure frequency, consistent with the results. If seizure severity is not influenced by CA2, or only indirectly, CNO would be unlikely to influence severity, again consistent with the results.

An important consideration is the degree to which DREADDs can be chronically activated when CNO is delivered in the drinking water for many weeks. Indeed, there is evidence that G-protein coupled receptors like DREADDs can desensitize (Gainetdinov et al., 2004; Urban and Roth, 2015; Black et al., 2016). However, the studies that report desensitization used injections, not drinking water, and all but one (Libbrecht et al., 2021) used hM4Di, not hM3Dq. There also are numerous reports that do not support the idea that DREADDs desensitize (Cheng et al., 2019; Page et al., 2019; Ewbank et al., 2020; Pozhidayeva et al., 2020; Xu et al., 2021). Moreover, if desensitization occurred in our study, the effects that we found would be underestimates. It should be noted, however, that in the epileptic brain desensitization may differ from the normal brain and future studies will be needed to address this possibility.

Another important point is that the metabolite of CNO, clozapine, increases the risk of seizures in humans (Wong and Delva, 2007; Williams and Park, 2015). Therefore, CNO might have increased the risk of seizures in our mice by metabolism to clozapine. However, we did not see effects of CNO in control mice, suggesting that the effects are indeed due to activation of CA2.

One of the limitations of the present study is that the pilocarpine model is only one model of TLE and the results may not generalize to other models of TLE or human TLE. On the other hand, this model had very robust seizures, lasting over 15 seconds and the vast majority of seizures were accompanied by easily recognized (stages 4-6) convulsive behavior. Seizures were frequent, providing statistical power to detect changes in seizure frequency.

There are several caveats related to the methods. Determination of viral expression provided good evidence of expression in CA2 but the measurements should be viewed only as estimates. One reason is that PCP4 may be an imperfect marker (Radzicki et al., 2023). There are also caveats of double-labeling: false double-labeling can occur if a red cell is on top of a green cell. However, we viewed both red and green channels to ensure a cell that was red was also green. Another caveat is that a section near the injection site was chosen for each animal, but viral expression will be lower as distance from that site increases. Therefore, the degree of viral expression for the entire dorsal hippocampus is likely to be lower than the expression we measured. The implication is that only a subset of dorsal CA2 expressed virus and the effects of chemogenetics on dorsal CA2 are likely to be an underestimation. Despite this caveat, Cre+ mice showed eDREADD-mCherry labeling in CA2 even after 6 weeks of vEEG, and none of the Cre-mice showed fluorescence, suggesting expression was specific for Cre+ mice and was stable.

Seizure analysis also has limitations. For example, detecting when a seizure begins and ends is not trivial. This limitation was not a major concern in this study because the majority of seizures began with a sentinel spike, something easy to define. Also, most seizures ended abruptly in all leads, making seizure termination readily defined. When seizures did not end in all electrodes at the same time, we selected the time when the first lead abruptly stopped its high amplitude spiking because it usually coincided with the abrupt end of the convulsive behavior. This was the same for all experimental groups. Another limitation is how to determine seizure frequency when seizures cluster, so we used a long recording period. In summary, we acknowledge the caveats and made efforts to limit them.

### Conclusions

This study showed that activating area CA2 using chemogenetics in the pilocarpine model exacerbated several aspects of epilepsy, such as seizure frequency, seizure duration, and the severity of clusters. Activation of CA2 also facilitated SE. Effects on seizure frequency were mainly detected in males. Taken together, there appears to be bidirectional regulation of seizures by CA2 using this epilepsy model. Given there are several unique marker genes for CA2, it may be possible to make use of genes specific to CA2 to tailor therapeutic strategy to inhibit CA2 in TLE in the future. Although the effects of CA2 inhibition (Whitebirch et al., 2022) or CA2 excitation (in the present study) are limited in magnitude, in both our studies there only was expression of DREADDs in a subset of CA2 neurons limited to dorsal hippocampus. A more widespread manipulation of CA2 activity, as would be possible using a CA2-specific pharmacological approach, may show a more effective control of seizure activity.

This study also sheds light on sex differences in the mouse pilocarpine model. Several analyses suggested that there were sex differences that make induction of SE, the severity of chronic seizures and the duration of convulsive seizures weaker in females than males. These sex differences may have contributed to the weaker effects of CA2 chemogenetic manipulation in females but more studies will be necessary to clarify mechanisms. Nevertheless, the results help address diverse reports in the literature of sex differences in epilepsy and provide new focus to this literature by pointing to a greater influence on certain aspects of seizures rather than all.

## Availability of data and materials

All data will be made available at www.isf.io upon acceptance. All materials are listed in the Methods with their vendor information.

## CRediT authorship contribution statement

Conceptualization: HES Methodology: JJL, MK, MR

Investigation: JJL, MK, MR

Formal analysis: JJL, HES

Writing – original draft – HES

Writing – review & editing – HES, BS, CPL, SAS

Funding acquisition – HES, SAS

## Consent for publication

All authors have approved the submission.

## Ethics in publishing

The authors have followed Elsevier’s publishing ethics policy (https://www.elsevier.com/about/policies-and-standards/publishing-ethics#4-duties-of-authors)

