## Supplemental material for "Chemogenetic activation of hippocampal area CA2 promotes acute and chronic seizures in a mouse model of epilepsy"

**SUPPLEMENTAL FIGURES**

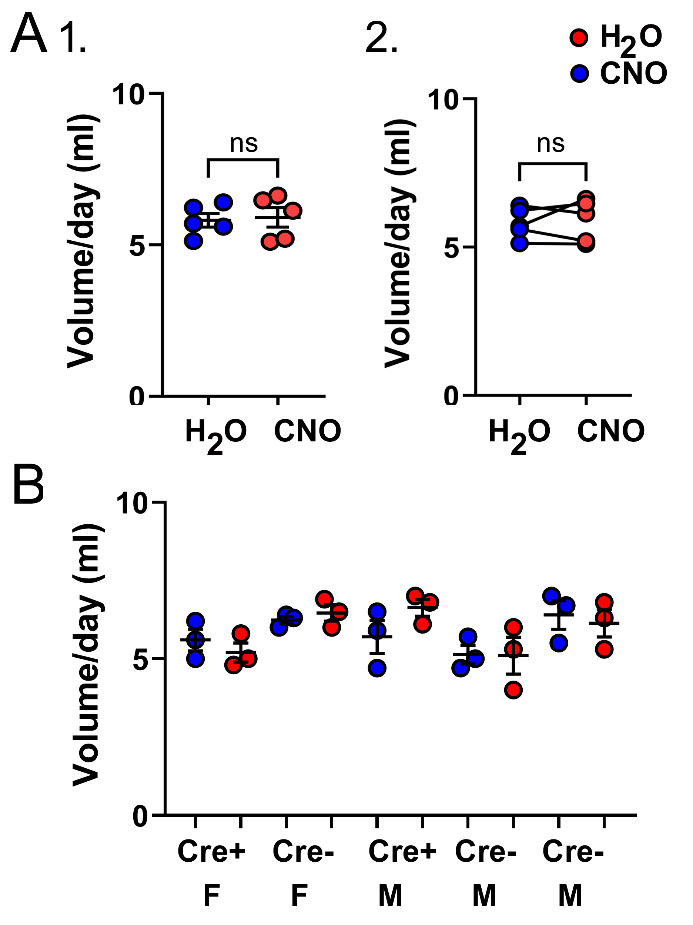

**Supplemental Figure 1. Volume of water consumed in mice treated with and without CNO.**

**A.** For 5 mice the mean volume of water consumed per day is shown for water with and without CNO. Differences were not significant (paired t-test, t=0.40, df=4, p=0.712). Red: H_2_O; blue: CNO.

1. Means ± SEM.

2. Paired comparisons.

**B.** All 5 mice are separated to show the measurements for water with and without CNO. Three measurements were made (1/week) for the 3 weeks treated with water and for water with CNO. Red: H_2_O; blue: CNO. F: female, M: male.

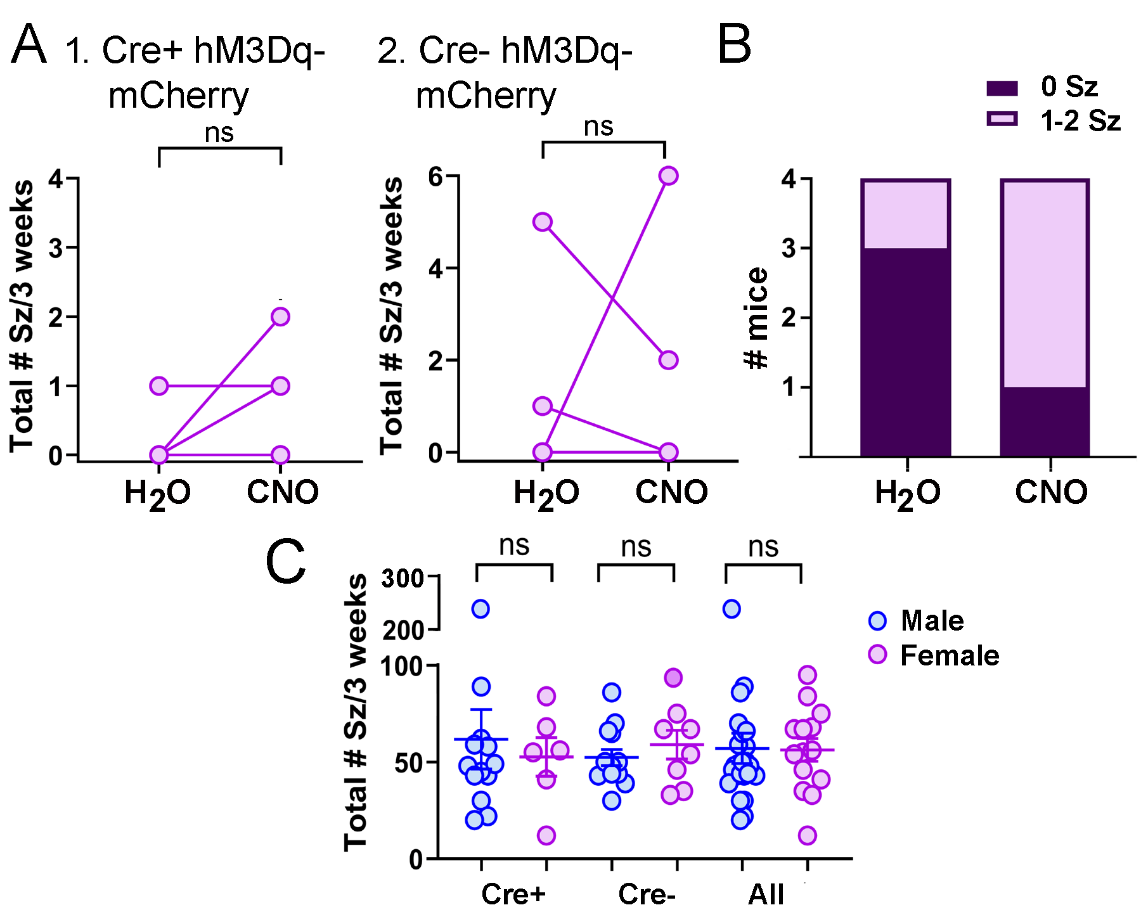
**Supplemental Figure 2.** Analysis of chronic seizures in female mice that were injected with pilocarpine but did not have SE.

**A.** Some female mice had pilocarpine but did not have SE, unlike males, which always had SE. Although it would not be expected that females without SE would develop chronic seizures, we tested them. Remarkably, some of these females had chronic seizures.

1. For the 4 Cre+ females without SE, seizures during the 3 weeks when mice drank water without CNO showed that there a seizure but only one mouse. In contrast, 3 of the same 4 mice had seizures when treated with CNO. However, the mean seizure frequency was not significantly affected by CNO (Wilcoxon matched-pairs signed rank test, W=3, p=0.500, n=4).

2. In 4 Cre- females without SE, differences in chronic seizure frequency while mice drank water with CNO vs. without CNO were not significant (Wilcoxon matched-pairs signed rank test, W=4, p>0.999, n=4; Fisher’s exact test, p>0.999, n=4).

**B.** The number of Cre+ mice without SE that had chronic seizures (pink) and did not have chronic seizures (purple) is shown for water with and without CNO. There were no significant differences (Fisher’s exact test, p=0.453).

**C.** Total # seizures for mice that had SE during treatment with normal water. Although some females did not have SE (A-B), those that had SE had a similar number of seizures as males. There were no differences between genotypes and no differences between pooled males and pooled females (Unpaired t-tests, Cre+ males vs. Cre+ females, t=0.38, df=17, p=0.706, 13 males, 6 females; Cre- males vs. Cre- females, t=0.84, df=19 p=0.414, 13 males, 8 females; all males vs. all females, t=0.07, df=38, p=0.943, 26 males, 14 females).
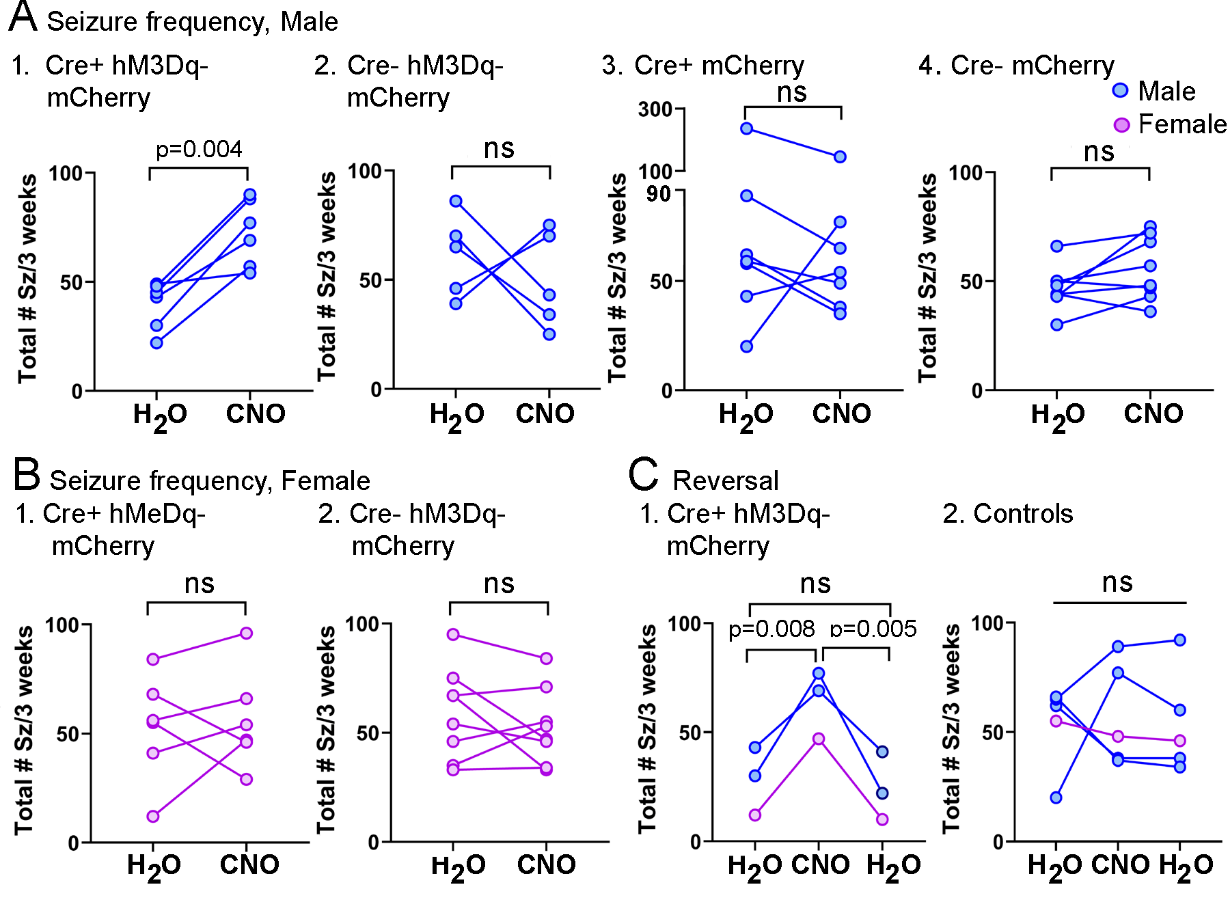

**Supplemental Figure 3.** Comparisons of seizure frequency in individual mice.

The data in Figure 1 are plotted with lines connecting the individual mice while drinking normal water or water with CNO.

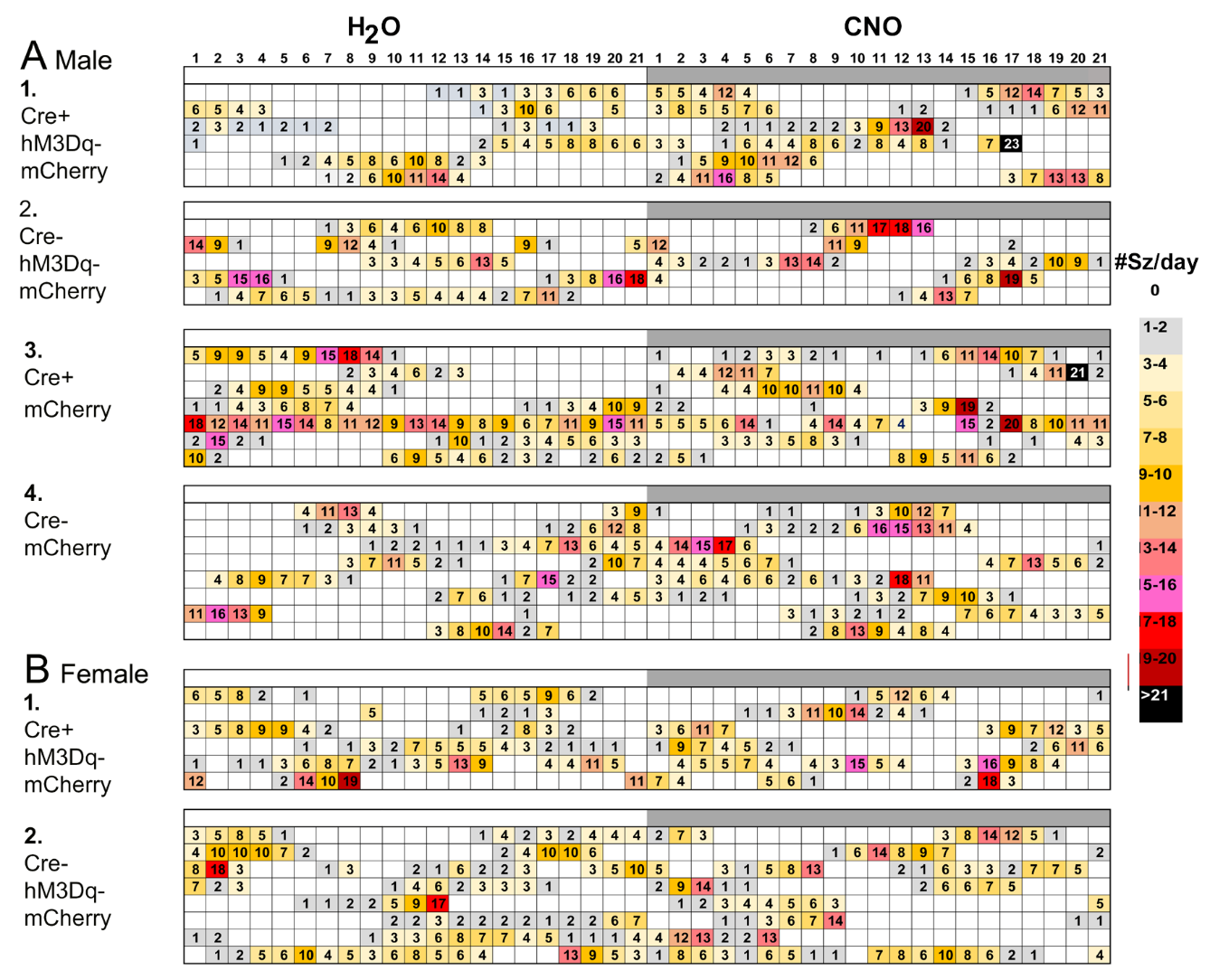
**Supplemental Figure 4.** Individual data for seizure frequency. Data and sample sizes are the same as for Figure 1.

**A.** For male mice, each experimental group is shown with the number of seizures/day noted for every day of the 6 week recording period. Each cell is a different day. The cells are color coded with darker colors reflecting more seizures/day (0, white; >21, black). The color code is shown to the right of the plots. Note that water treatment is on the left and CNO treatment on the right to optimize comparisons, but some animals were treated with CNO first and others treated with CNO second.

1. Cre+ mice injected with AAV-hM3Dq-mCherry. CNO treatment days included more color, supporting the data in Figure 1 suggesting that more seizures occurred.

2. Cre- mice injected with AAV-hM3Dq-mCherry.

3. Cre+ mice injected with AAV-mCherry.

4. Cre- mice injected with AAV-mCherry.

**B.** Data for female mice.

1. Cre+ mice injected with AAV-hM3Dq-mCherry. Consistent with the lack of a significant effect of CNO on seizure frequency in Figure 1, colors were similar for both the treatment with and without CNO.

2. Cre- mice injected with AAV-hM3Dq-mCherry.

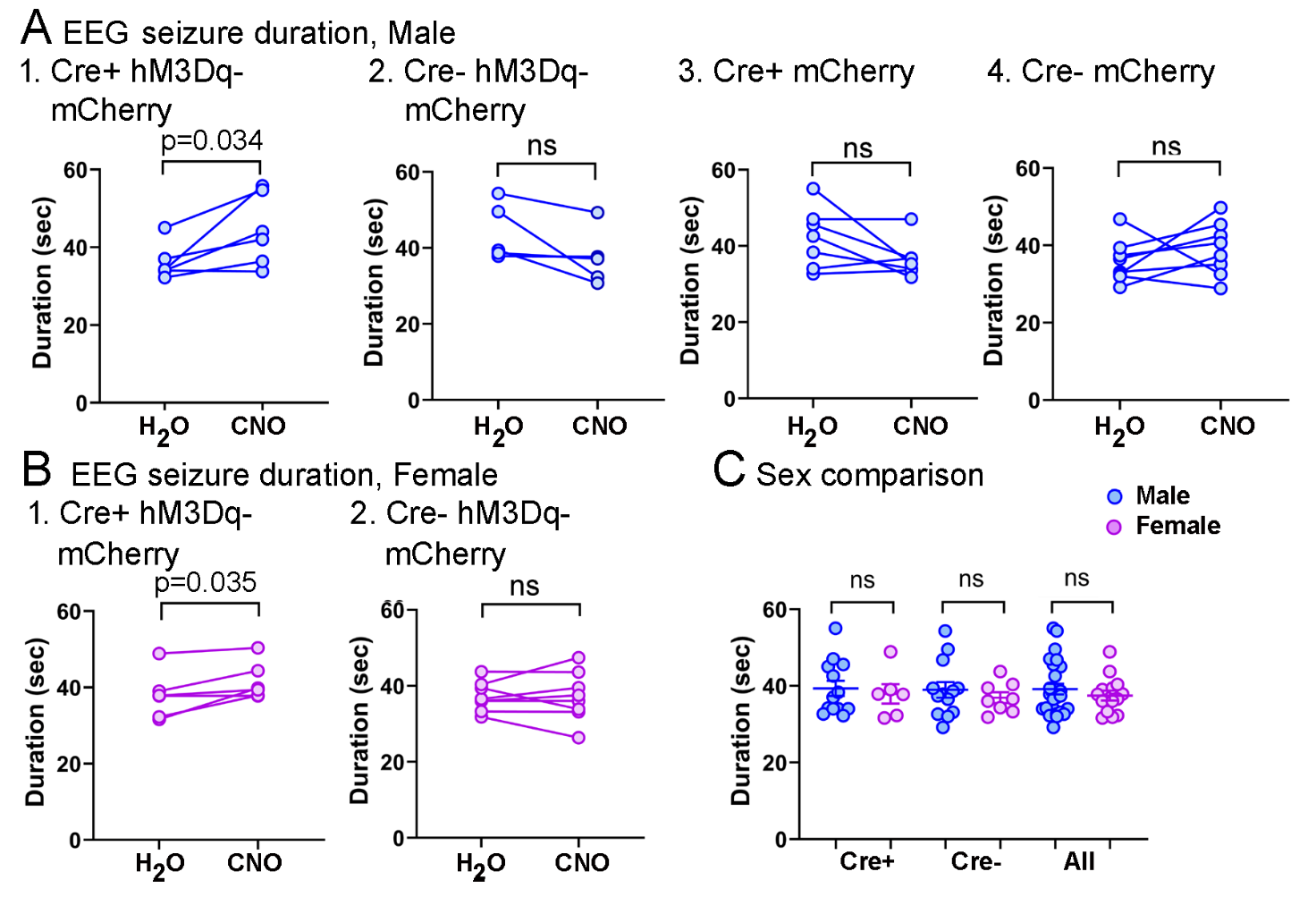
**Supplemental Figure 5. Comparisons of EEG seizure duration in individual mice.**

**A-B.** The data from Figure 2 are plotted with lines connecting individual mice treated with normal water or water with CNO.

**C.** There were no sex differences in the duration of EEG seizures (unpaired t-tests, Cre+ males vs. Cre+ females, t=0.43, df=17, p=0.672, n=13 males, 6 females; Cre- males vs. Cre- females, t=0.72, df=19, p=0.480, n=13 males, 8 females; all males vs. all females, t=0.80, df=38, p=0.428, n=26 males, 14 females).

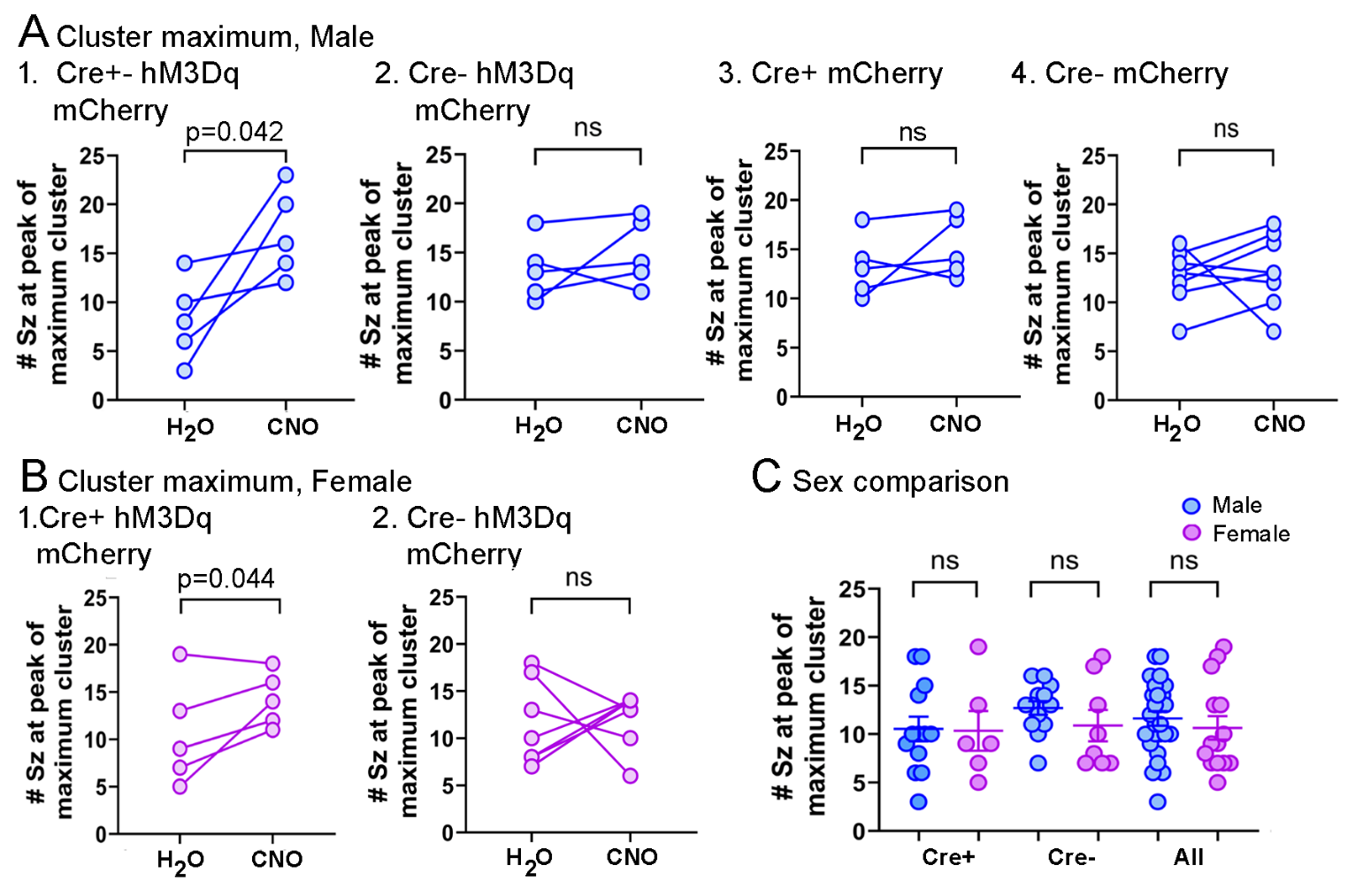
**Supplemental Figure 6. Comparisons of the peak of seizure clusters in individual mice.**

**A-B.** The data from Figure 3 are plotted, showing the data for each mouse.

**C.** There were no sex differences in the peak of seizure clusters (unpaired t-tests, Cre+ males vs. Cre+ females, t=0.09, df=17, p=0.931, n=13 males, 6 females; Cre- males vs. Cre- females, t=1.17, df=19, p=0.256, n=13 males, 8 females; all males vs. all females, t=0.72, df=38, p=0.479, n=26 males, 14 females).

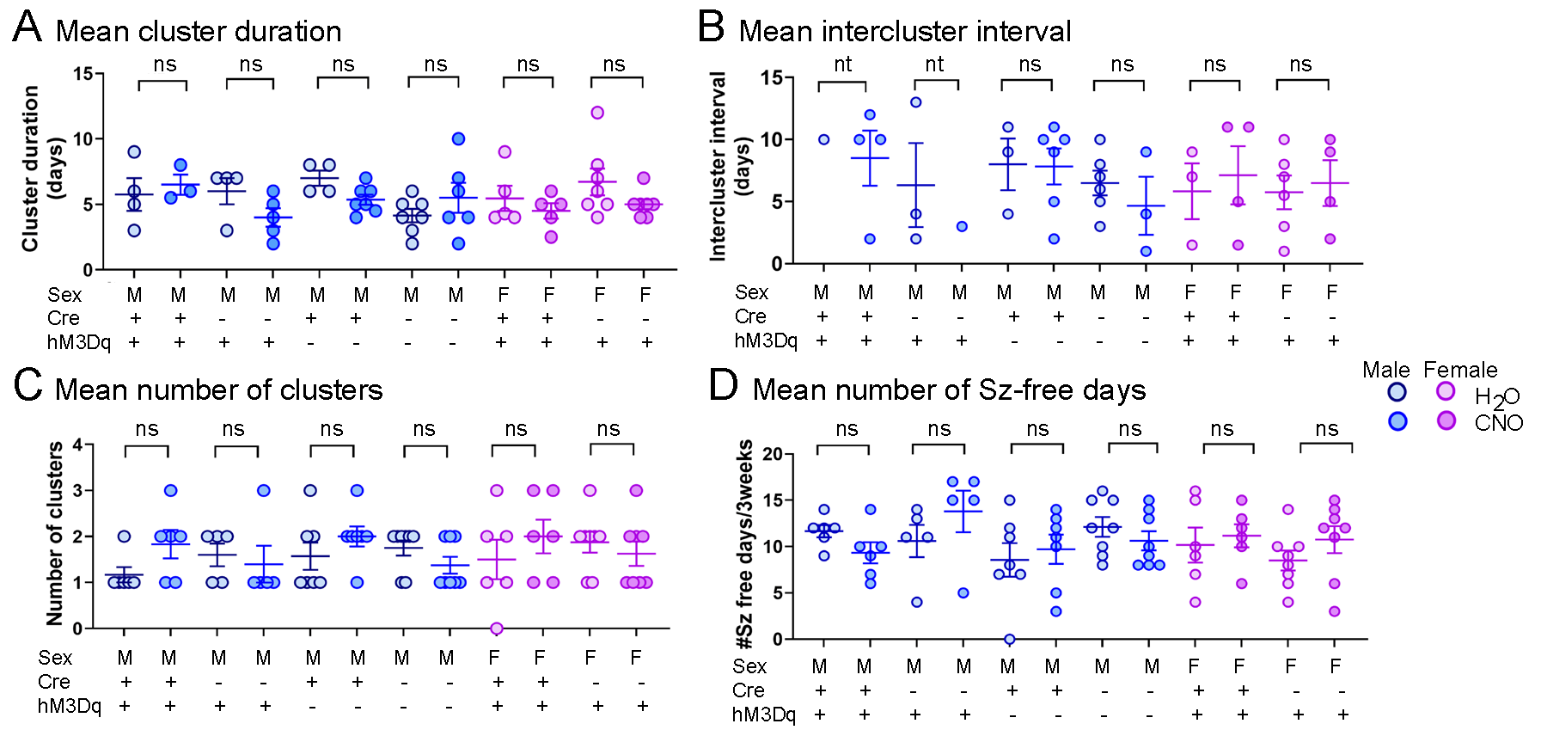

**Supplemental Figure 7. Measurements of seizure clusters that did not show effects of CNO.**

**A.** The mean cluster duration was measured for each animal during treatment with normal water and water containing CNO. In many animals the cluster duration was not possible to measure because the 3 week-long recording started and ended during a cluster. Therefore, data were analyzed with unpaired t-tests. For A-D there were no significant differences, but the sample sizes were often low so the results should be considered exploratory.

**B.** There were no significant differences in the mean intercluster interval. In some animals the interval was unclear because the 3 weeks of video-EEG started and ended during a cluster. Therefore, statistics were conducted using unpaired t-tests. In groups with fewer than 3 mice no statistical comparisons were possible. There were no significant differences in those groups with >3 mice. nt=not tested because there was only 1 cluster, so it was not possible to determine the intercluster interval.

**C.** The number of clusters in 3 weeks was compared. There were no statistical differences.

**D.** There were no significant differences in the number of seizure-free days per 3 weeks. For A-D, the statistical comparisons are in Supplemental Table 4.

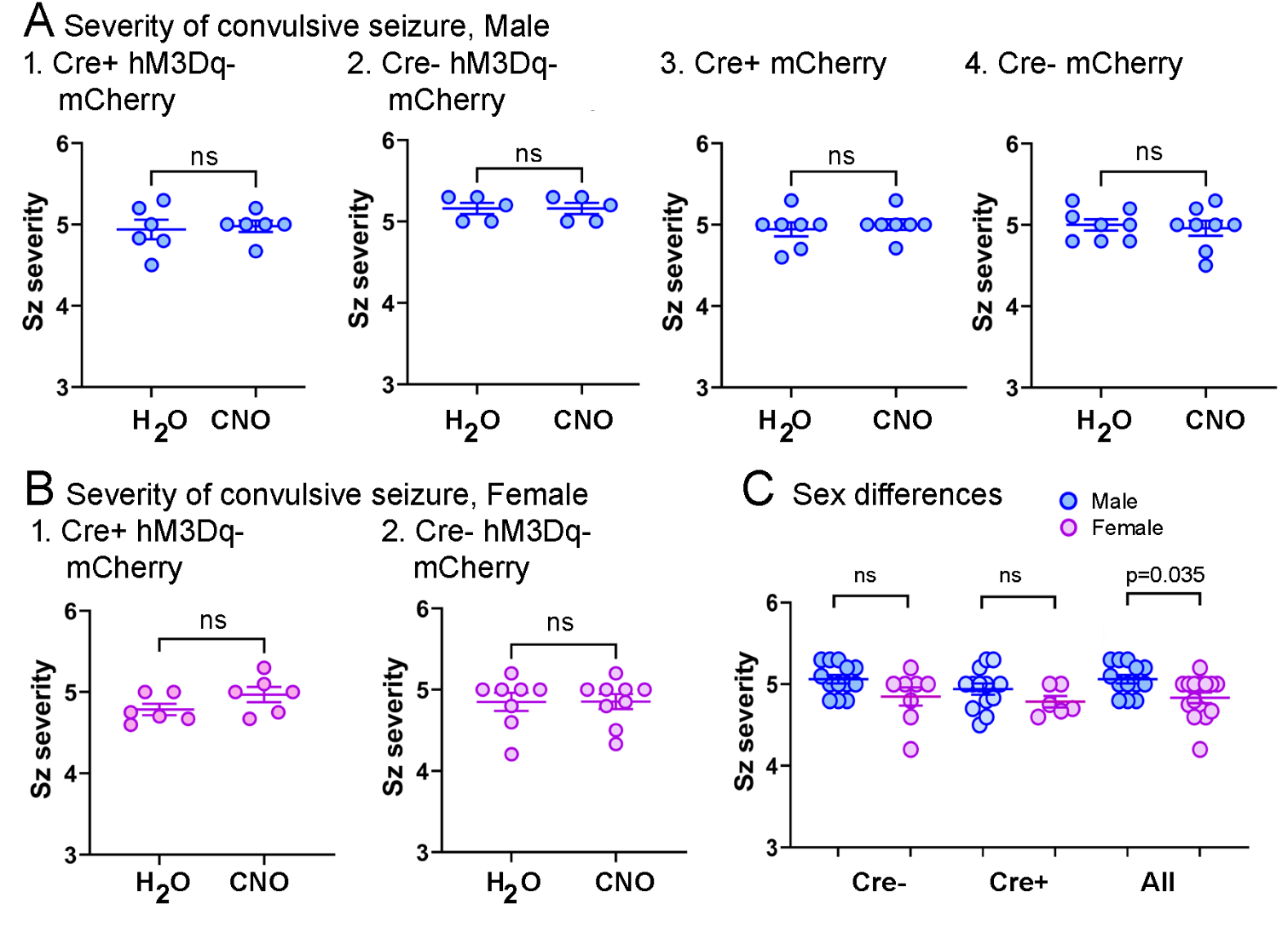

**Supplemental Figure 8.** Severity of convulsive seizures.

**A.** Severity of convulsive seizures showed no effects of CNO in males.

Wilcoxon tests: A1, W=2.00, p=0.875, n=6; A2, W=0, p>0.999, n=5; A3, W=4.00, p=0.625, n=7; A4, W=-4.00, p=0.812, n=8.

**B.** Severity of convulsive seizures did not show effects of CNO in females.

Wilcoxon tests: B1, W=6.00, p=0.250, n=6; B2, W=-6.00, p=0.500, n=8.

**C.** There were sex differences in the severity of seizures. In comparisons of the seizures during the treatment with normal water, there was no effect of sex in Cre- or Cre+ mice. However, when genotypes were pooled, females had less severe seizures (Mann-Whitney *U*-tests, Cre- males vs. Cre- females, *U*=30.5, p=0.114, n=13 males, 8 females; Cre+ males vs. Cre+ females, *U*=24, p=0.189, n=13 males, 6 females; all males vs all females, *U*=110, p=0.035, n=26 males, 14 females).

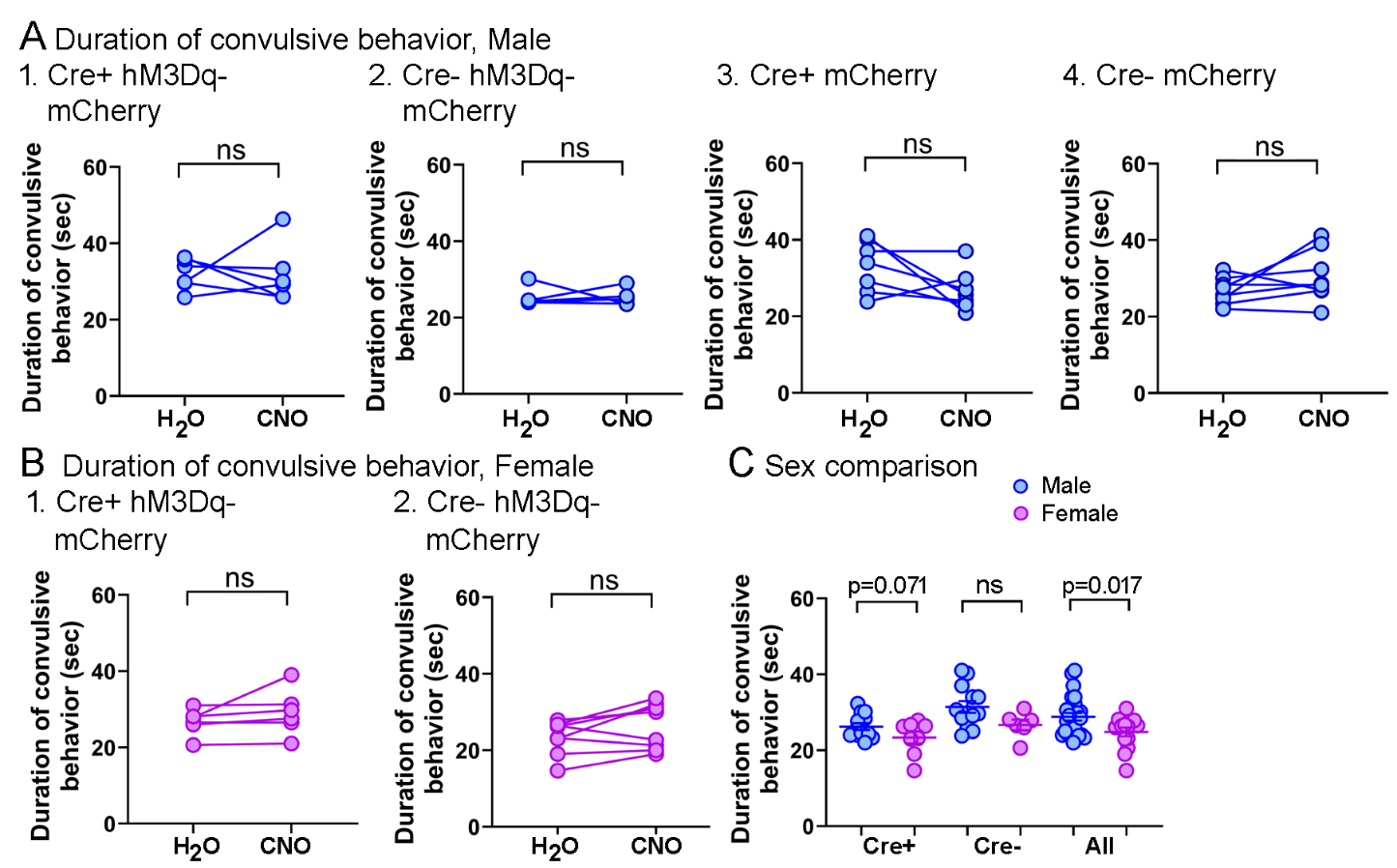

**Supplemental Figure 9. Comparisons of the duration of convulsive behavior.**

**A.** The duration of convulsive behavior during chronic seizures was measured in 10-15 seizures/mouse, selected at random. There was no effect of CNO in males. Paired t-tests: A1, t=0.83, df=5, p=0.443, n=6; A2, t=0.03, df=4, p=0.978, n=5; A3, t=1.04, df=6, p=0.338, n=7; A4, t=1.53, df=7, p=0.169, n=8.

**B.** The duration of convulsive behavior did not show effects of CNO in females. Paired t-tests: B1, t=1.44, df=5, p=0.210, n=6; B2, t=1.84, df=7, p=0.109, n=8.

**C.** There were sex differences in the duration of convulsive behavior. In comparisons of the seizures during the treatment with normal water, there was no effect of sex in Cre- or Cre+ mice. However, when genotypes were pooled, females had shorter convulsive behavior during seizures (unpaired t-tests: Cre+ males vs. Cre+ females, t=1.92, df=17, p=0.071, n=13 males, 6 females; Cre- males vs. Cre- females, t=1.70, df=19, p=0.105, n=14 males, 8 females; all males vs. all females, t=2.50, df=38, p=0.017, n=26 males, 14 females).

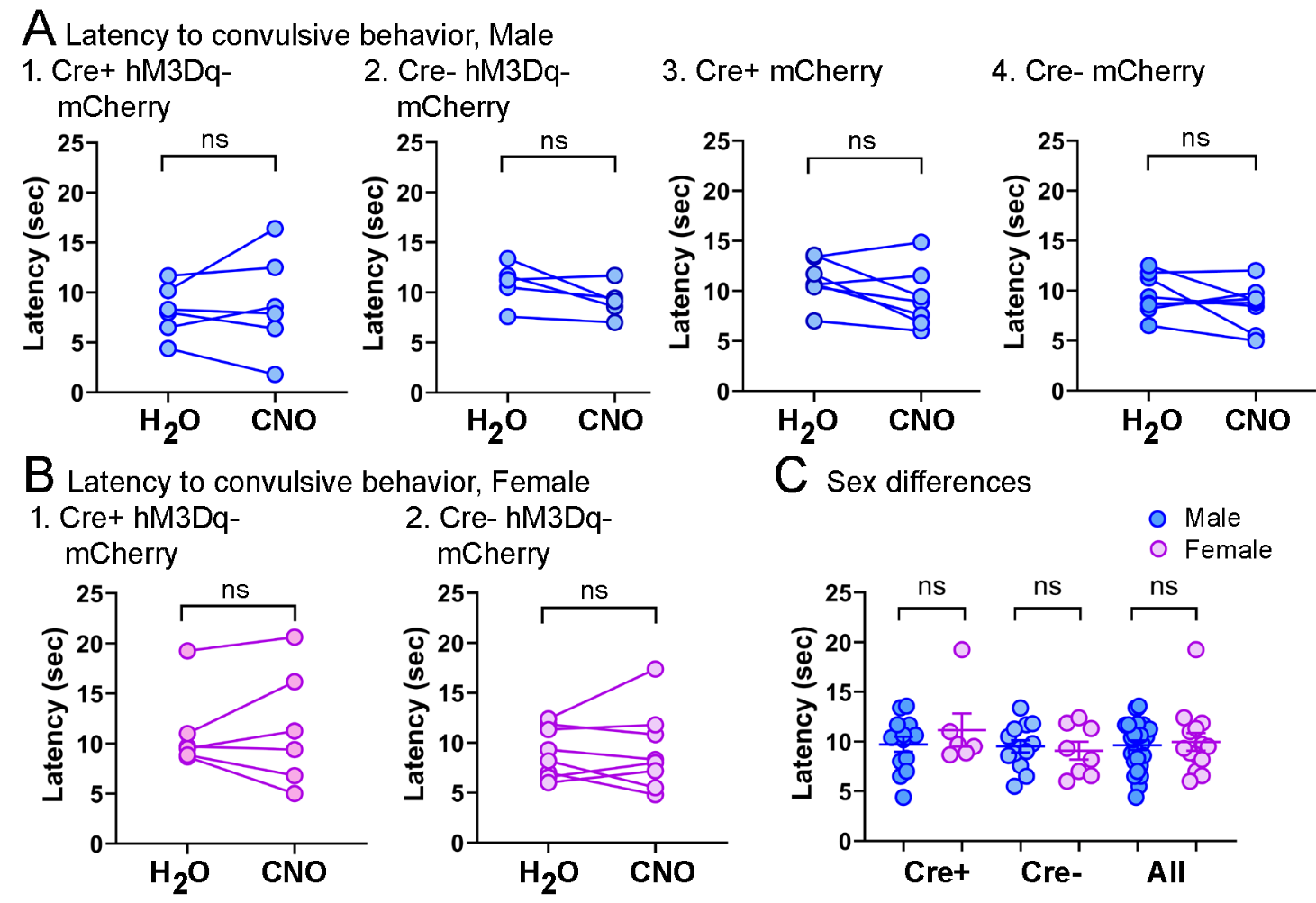
**Supplemental Figure 10. Comparisons of the latency to convulsive behavior during a seizure**

**A.** The latency from the start of the seizure, defined by EEG, to the latency of the convulsive behavior, defined by video, showed no effect of CNO in males. Paired t-tests: A1, t=0.58, df=5, p=0.587, n=6; A2, t=2.00, df=4, p=0.117, n=5; A3, t=1.92, df=6, p=0.104, df=7; A4, t=0.78, df=7, p=0.461, df=8.

**B.** The latencies to convulsive behavior did not show effect of CNO in females. Paired t-tests: B1, t=1.94, df=4, p=0.125, n=5; B2, t=0.16, df=7, p=0.874, n=8.

**C.** There were no sex differences in the latencies. In comparisons of the seizures during the treatment with normal water, there was no effect of sex in Cre- or Cre+ mice or when genotypes were pooled (unpaired t-tests, Cre+ males vs. Cre+ females, t=0.14, df=16, p=0.889, n=13 males, 6 females; Cre- males vs. Cre- females, t=0.29, df=16, p=0.779, n=13 males, 5 females; all males vs. all females, t=0.20, df=33, p=0.842, n=26 males, 10 females).

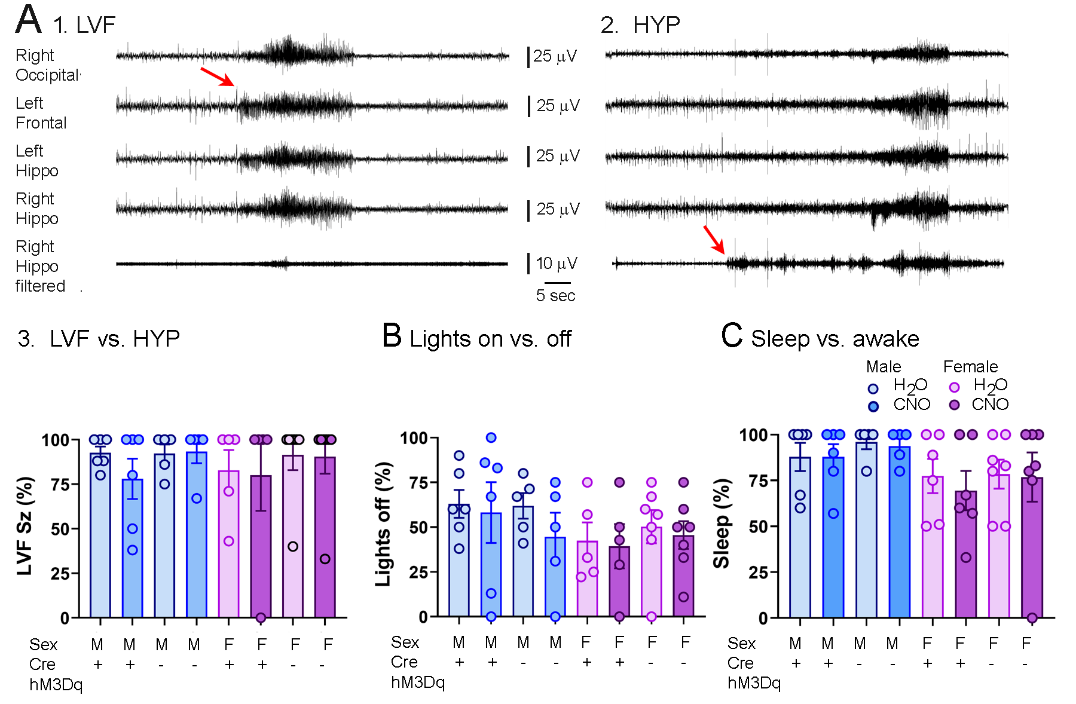
**Supplemental Figure 11.** The type of seizure and when they occurred did not show an effect of CNO.

**A.**

1. An example of a seizure referred to as low-voltage fast (LVF). There is a sentinel spike at the start (arrow) followed after a delay by high frequency, high amplitude activity. At the bottom is the record from the right hippocampus, filtered to show high frequencies (>80 Hz, FIR filter, high pass, Spike2 software).

2. An example of a seizure with an onset that is hypersychronous (HYP). There is no clear sentinel spike. At the bottom is the record from the right hippocampus, filtered as for the LVF seizure but showing much more high frequency activity at the start of the seizure. The onset of this activity defined seizure onset (arrow).

3. Ten seizures were selected at random for each animal. The percent that were LVF is shown. There were no significant differences in the % of seizures that were LVF between normal water and water with CNO.

**B.** There were no detectable effects of CNO on the % of seizures occurring during sleep.

**C.** There were no detectable effects of CNO on the % of seizures occurring when lights were off vs. lights on. Statistical information for A3, B, and C are in Supplemental Table 5.

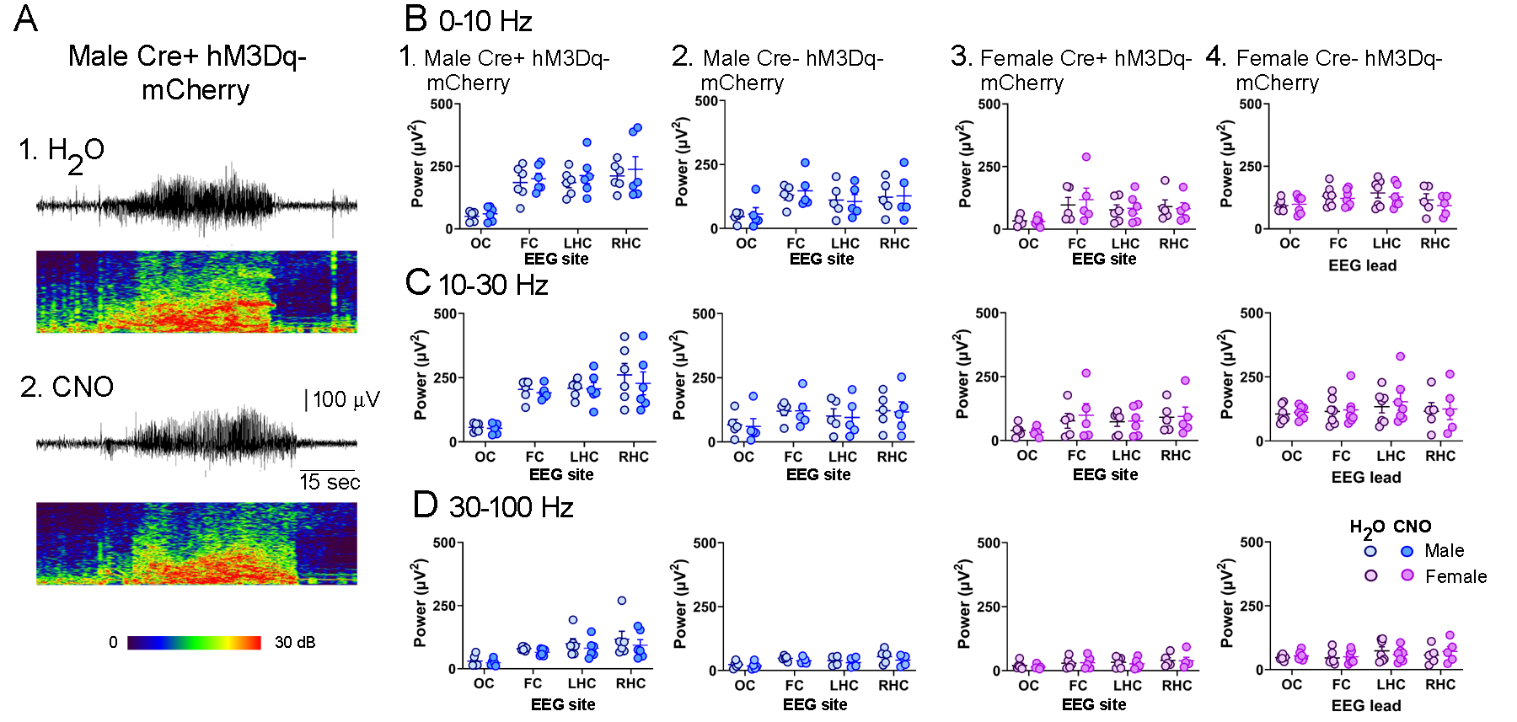

**Supplemental Figure 12.** Power of chronic seizures was unaffected by CNO.

**A.** Seizures in a male Cre+ injected with AAV-hM3Dq-mCherry.

1. An example of a seizure while a mouse was treated with normal water. The EEG trace from the right hippocampal electrode is shown above the spectrogram.

An example of a seizure in the same mouse during the treatment with CNO. The recording is from the right hippocampal electrode.

B. Comparisons of power in the 0-10 Hz band (mainly reflecting delta and theta rhythms) while mice received regular water or CNO in their water. Statistical comparisons are in Supplemental Table 6. Note that one female Cre- mouse could not be used because all electrodes had artifacts during seizures.
1. Male Cre+ mice injected with AAV-hM3Dq-mCherry.
2. Male Cre- mice injected with AAV-hM3Dq-mCherry.
3. Female Cre+ mice injected with AAV-hM3Dq-mCherry.
4. Female Cre- mice injected with AAV-hM3Dq-mCherry.
C. Comparisons of power in the 10-30 Hz band (including mu and beta rhythms).
D. Comparisons of power in the 30-100 Hz band (including gamma).

**SUPPLEMENTAL TABLES**

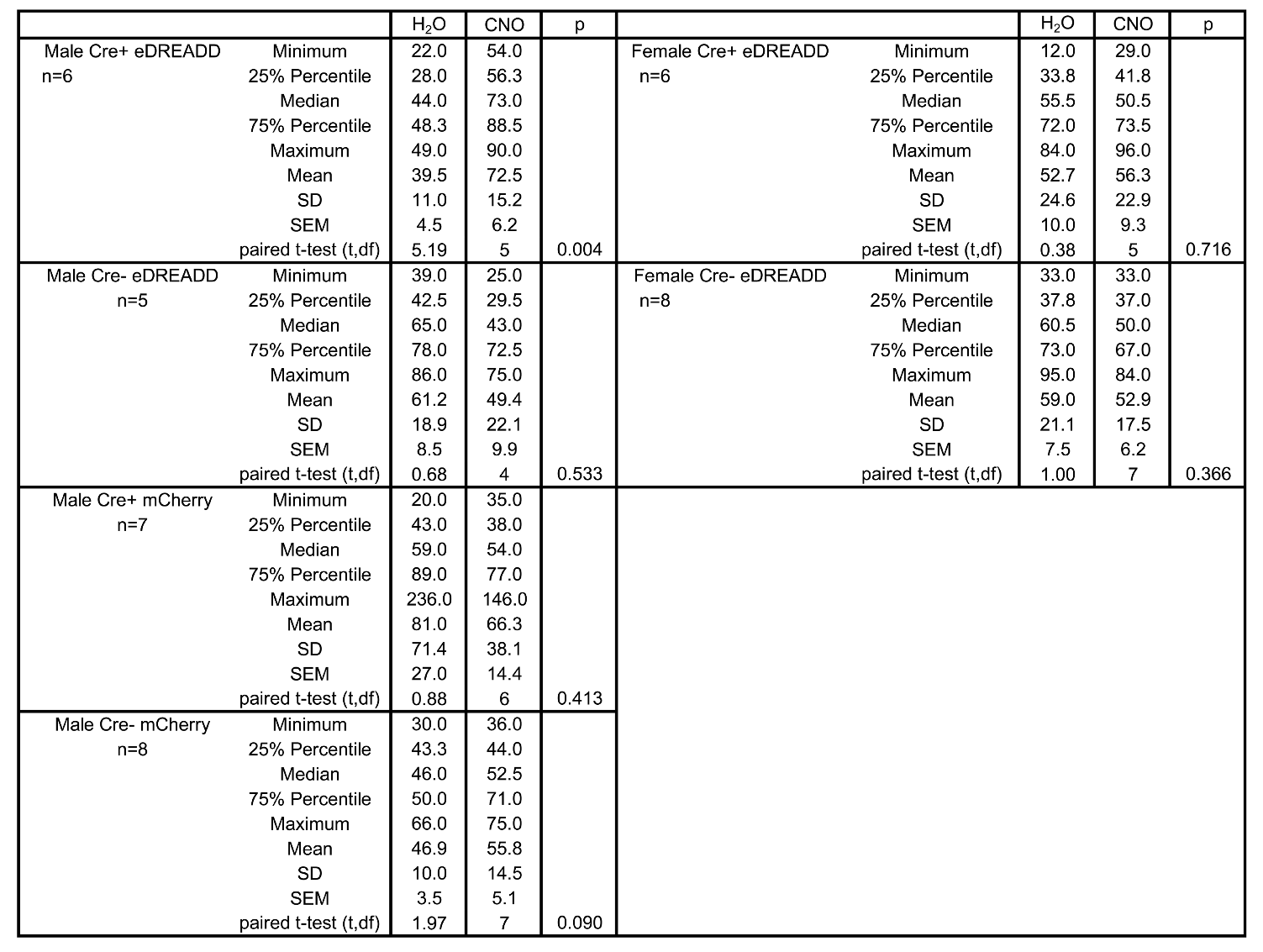

**Supplemental Table 1. Comparisons of seizure frequency.**

The statistical comparisons for Figure 1 are shown, as well as measurements of the minimum, median, maximum, 25%, and 75% percentiles.

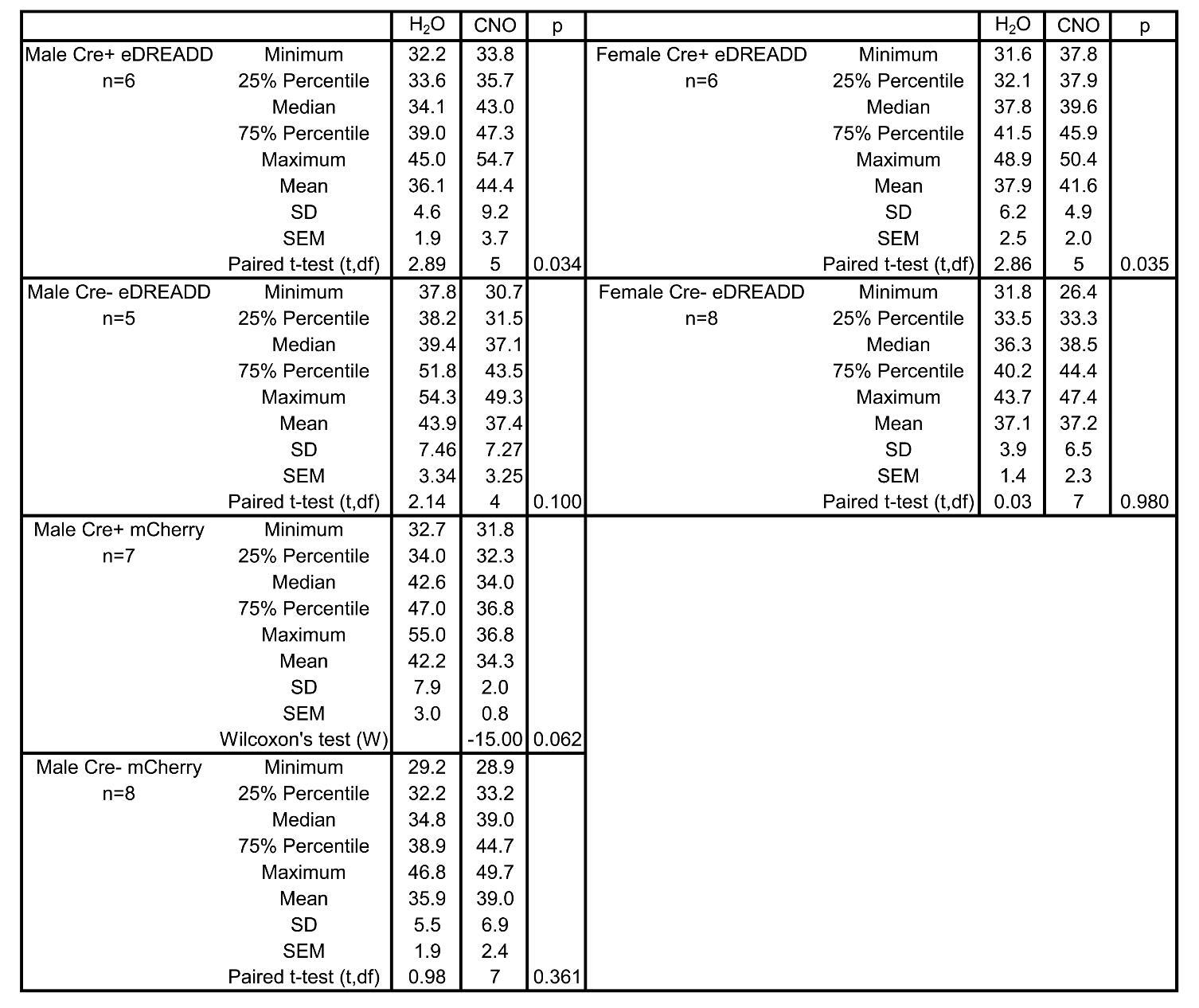

**Supplemental Table 2. Comparisons of EEG seizure duration.**

The statistical comparisons for Figure 2 are shown, as well as measurements of the minimum, median, maximum, 25%, and 75% percentiles. Note that the Cre+ males injected with AAV-hM3Dq-mCherry the variances in the means when mice drank water without CNO vs. water with CNO were unequal. Therefore, data were log transformed to resolve the heteroscedasticity. The means, medians, maximums and 25/75 percentiles of the raw data are shown. The details of the paired t-test using transformed data are also shown. Note that the p value was significant for the transformed data (p=0.034) and prior to transformation (p=0.041).

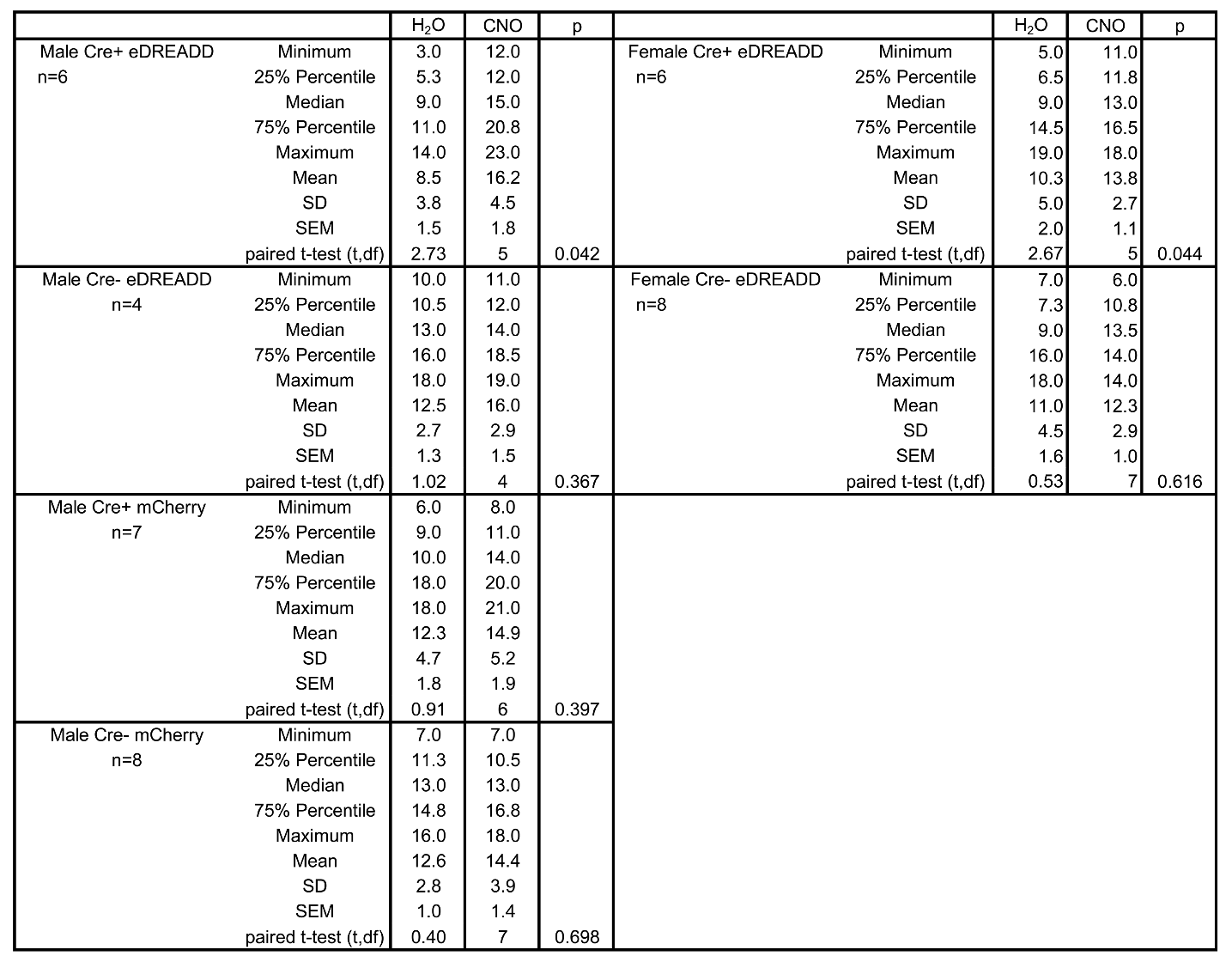

**Supplemental Table 3. Comparisons of the peak of seizure clusters.**

The statistical comparisons for Figure 3 are shown, as well as measurements of the minimum, median, maximum, 25%, and 75% percentiles.

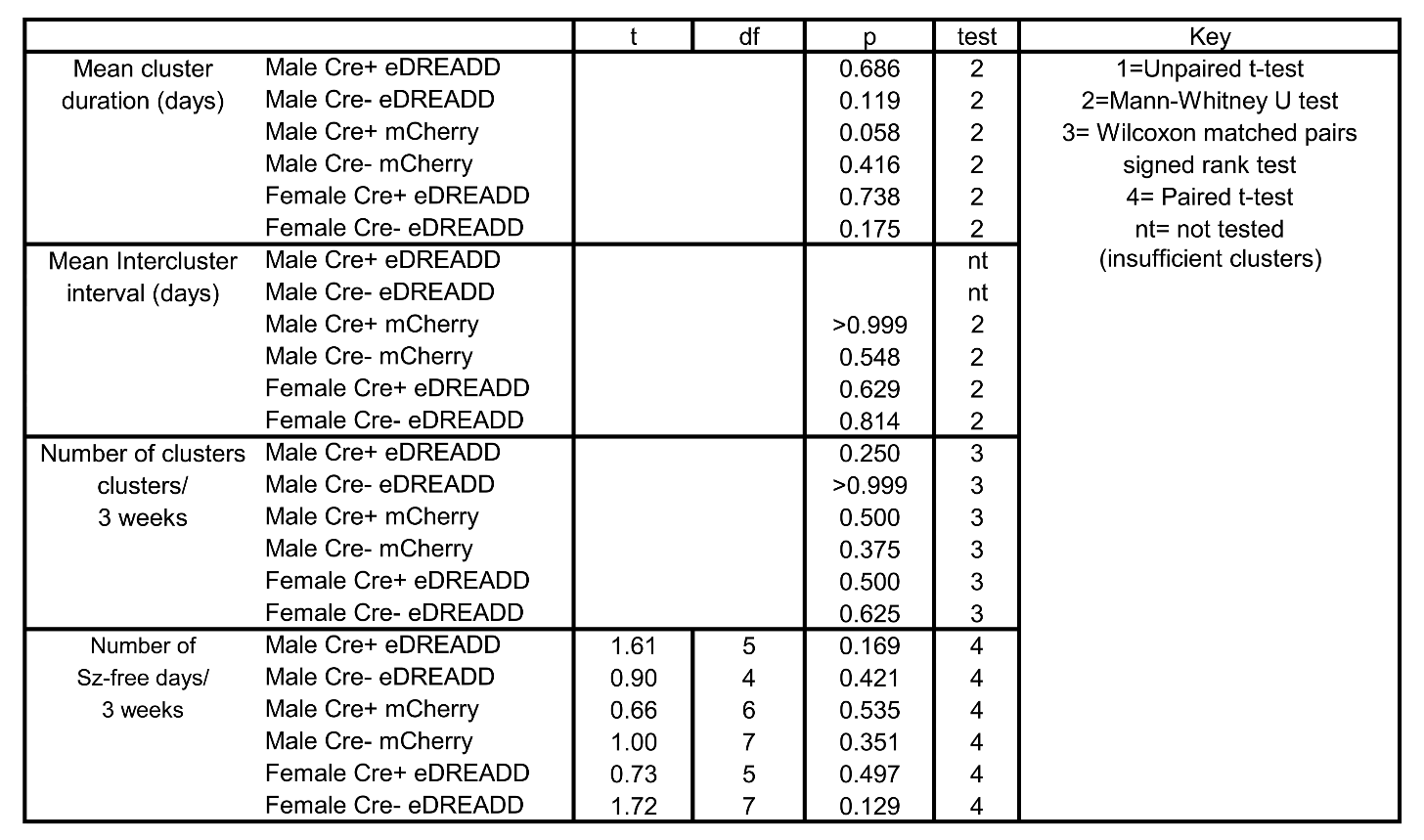
**Supplemental Table 4. Additional characteristics of seizure clusters.**

The statistical comparisons for cluster duration, intercluster interval, the numbers of clusters and the number of Sz-free days/3 weeks are shown. The statistical tests were U= Unpaired t-test (when some mice had values only for water or only for CNO); P= Paired t-test (for mice that had data during water and water with CNO); W= Wilcoxon signed-rank test (same as the paired t-test except data were not parametric). nt reflects that no statistical comparisons were made because the measurement of intercluster interval required at least 2 clusters and some mice lacked 2 clusters.

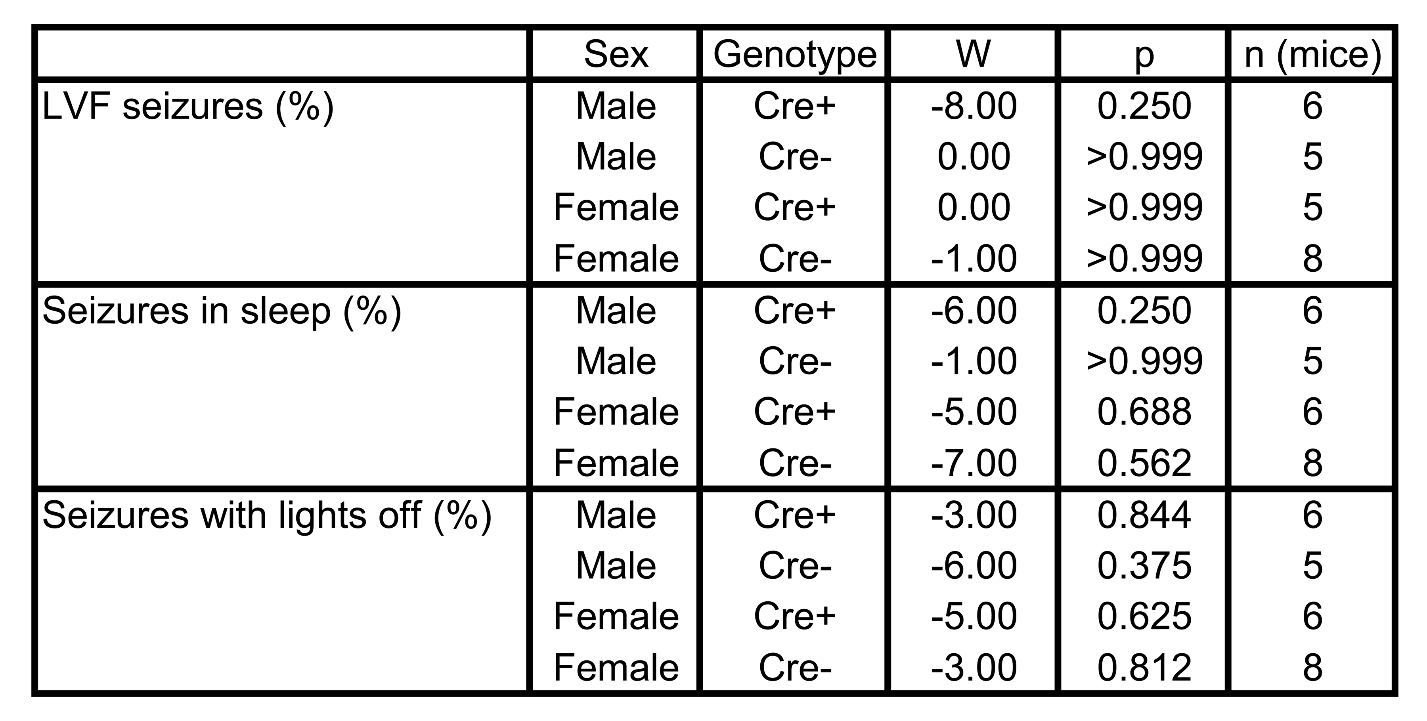

**Supplemental Table 5. Additional measurements of chronic seizures.**

Seizures were characterized as having a LVF or HYP onset. LVF seizures (%) is the percent of all LVF and HYP seizures that were LVF in their seizure onset pattern. Seizures in sleep (%) is the percent of all seizures that occurred during sleep. Seizures with lights off (%) is the percent of all seizures that occurred when lights were off. Statistics were Wilcoxon signed-ranks test.

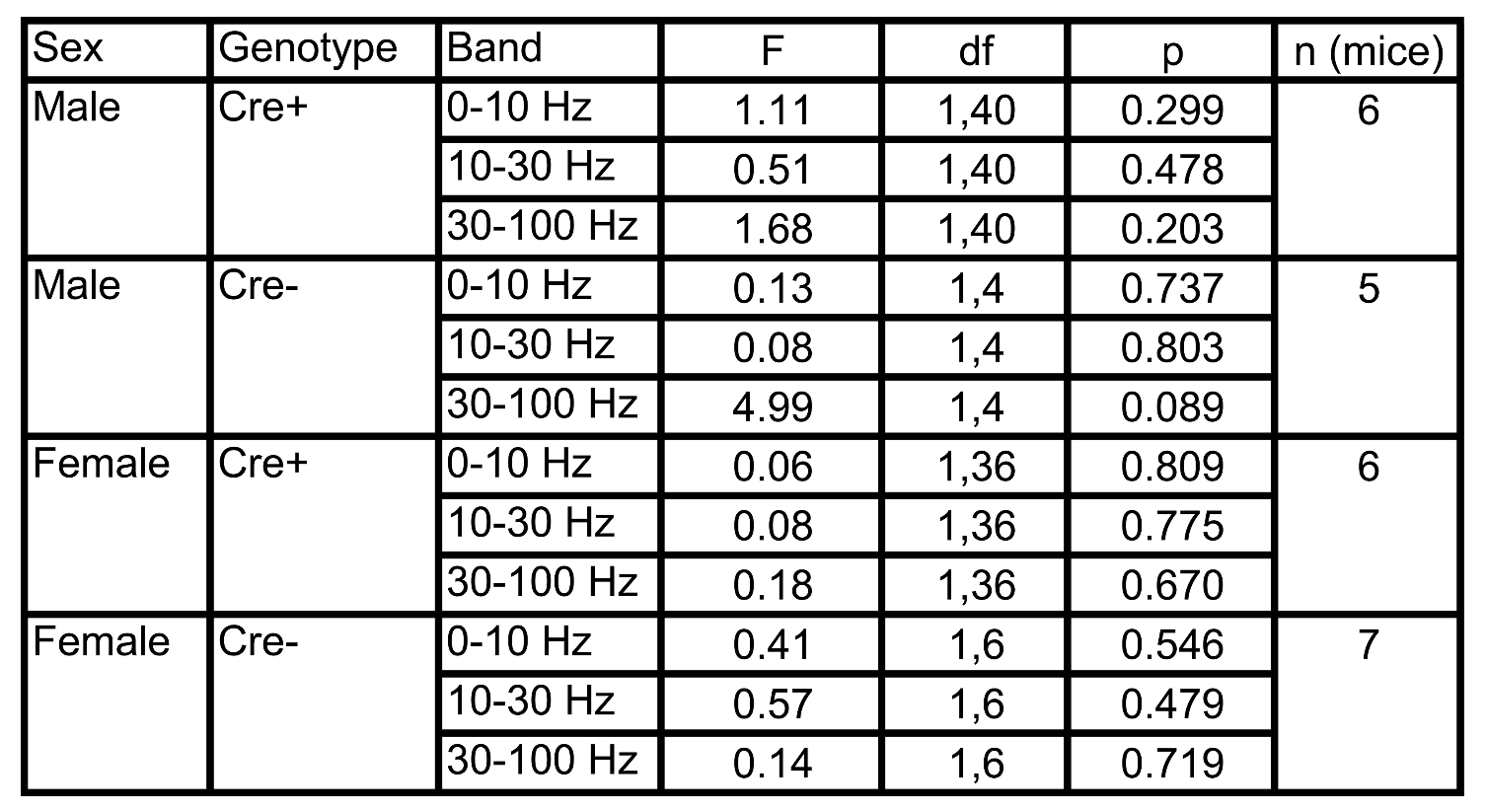

**Supplemental Table 6. Statistical comparisons of power.**

Comparisons of power were made for Cre+ and Cre- mice injected with AAV-hM3Dq-mCherry. Power was quantified for 0-10, 10-30, and 30-100 Hz frequency bands. Calculations were made for each of the four electrodes in each mouse for 5 seizures during water and 5 seizures during water with CNO (see Methods). Statistical comparisons of the seizures during treatment with water and CNO used two-way ANOVAs with the type of treatment (water or water with CNO) and electrode location as factors. However, Cre- data used a mixed effects analysis because some electrodes had artifacts during seizures and could not be used. Note that one female Cre- mouse could not be used because all electrodes had artifacts during seizures.
